# *Candida spp.* suppress neutrophil reactive nitrogen species to evade killing

**DOI:** 10.1101/2023.11.02.565128

**Authors:** Thomas B. Burgess, Ffion R. Hammond, Piotr T. Szkuta, Amy Lewis, Stella Christou, Kieran A Bowden, Tihana Bicanic, Lynne R Prince, Kathryn R. Ayscough, David G. Partridge, Simon A. Johnston, Alison M. Condliffe, Philip M. Elks

## Abstract

*Candida albicans* is a human commensal that can cause life-threatening invasive infection in immunocompromised individuals. Human immunity to *C. albicans* infection is thought to be largely dependent on neutrophil reactive oxygen and nitrogen species (ROS/RNS) generation by neutrophils. Despite this, our understanding of innate immune killing and escape by *C. albicans* is primarily studied in macrophages and the precise mechanisms of evasion are unclear in neutrophils. Here we sought to determine the importance of neutrophil reactive nitrogen species (RNS) production during *C. albicans* infection *in vivo*. Using a zebrafish model, we found that *C. albicans* rapidly downregulated neutrophil RNS below basal levels during the first day post infection, a time at which neutrophil RNS is upregulated in bacterial infections as an important host-defense mechanism, indicating fungal evasion of host neutrophils. We confirmed downregulation of RNS in human primary neutrophils and with clinical *Candida* spp. isolates, including emerging human pathogens *C. auris* and *C. glabrata*. Using a zebrafish *arginase 2* transgenic line and a *C. albicans car1*Δ mutant, we show that both host and fungal arginase contribute the reduction in neutrophil RNS. Despite pathogen downregulation, upregulation of neutrophil RNS via Hypoxia inducible factor (Hif)-1α stabilisation, was sufficient to improve *C. albicans* infection survival, dependent on the presence of neutrophils and Nitric oxide synthase 2 (Nos2). Finally, restoration of neutrophil RNS, via Hif-1α stabilisation, was synergistic with clinically relevant antifungal treatment, increasing survival and clearance of *C. albicans* infections. Together, these data demonstrate that restoration of the neutrophil RNS response in *C. albicans* infection improves infection outcomes, highlighting the potential of targeting Hif-1α and RNS in host directed therapies against fungal infections.

## Introduction

*Candida albicans* (*C. albicans*) is a commensal fungus, colonising 70% of healthy individuals (Witherden *et al*., 2017) and can cause superficial mucosal infections with negligible mortality. However, *C. albicans* can act as an opportunistic pathogen capable of causing life-threatening, invasive disease in immunocompromised individuals (Witherden *et al*., 2017; Flores-Maldonado *et al*., 2019; Pellon, Sadeghi Nasab and Moyes, 2020). There are approximately 750,000 cases of severe candidiasis worldwide per year and the estimated mortality rate is high, estimated to be around 50% (Bongomin *et al*., 2017; WHO, 2022), likely an underestimation due to poor surveillance mechanisms worldwide (Lamoth *et al*., 2018; Almeida, Rodrigues and Coelho, 2019). This high mortality, combined with a lack of fungal vaccines and rising rates of antifungal resistance, has resulted in *C. albicans* being designated as one of four critical priority fungal pathogens by the World Health Organization, with another *Candida* spp., the emerging human pathogen *Candida auris*, also a critical priority pathogen (WHO, 2022). Hence, urgent research into new therapies is required.

*C. albicans* is detected by pattern recognition receptors, including Toll-like receptors, C-type lectin receptors and the Rig-I like receptor MDA-5, resulting in a downstream pro-inflammatory response and innate immune cell recruitment (Burgess, Condliffe and Elks, 2022). Although many immune cell types are recruited to the sites of *C. albicans* infection, neutrophils are one of the most potent killers of *C. albicans*, with neutropenic patients being particularly at risk of invasive disease (Uzun *et al*., 2001; Brown, 2011). Following recruitment, neutrophils have multiple mechanisms of eliminating *C. albicans*, including the production of reactive oxygen (ROS) and nitrogen (RNS) species (Winterbourn *et al*., 2006). Pharmacological inhibition of nitric oxide synthase (NOS) leads to increased mortality in *C. albicans* infections in mice *in vivo* and reduced candidacidal activity in murine macrophages and peritoneal cells *in vitro*, showing the importance of RNS-mediated fungicidal activity (Rementería, García-Tobalina and Sevilla, 1995; Vazquez-Torres *et al*., 1995; Vazquez-Torres, Jones-Carson and Balish, 1996; Navarathna, Lionakis and Roberts, 2019). *C. albicans’* ability to rapidly adapt to changes in environmental pH, metabolic flexibility and a robust stress response allows colonisation of a wide range of niches within the host (Mayer, Wilson and Hube, 2013). As part of these mechanisms, *C. albicans* is able to subvert the host innate immune response, including ROS and RNS (Luo *et al*., 2013). Despite *Candida* spp. being primarily controlled by neutrophils in human disease, most immune evasion studies have used murine macrophage cells. This is, in part, because neutrophils are a more challenging cell type to study *in vitro* due to their short lifespan and ease of accidental activation. *C. albicans* can suppress ROS and RNS production by macrophages *in vitro* (Wellington, Dolan and Krysan, 2009; Collette, Zhou and Lorenz, 2014). Culture supernatant also suppressed macrophage RNS to a lower degree implying secreted compounds are partially involved in RNS suppression (Collette, Zhou and Lorenz, 2014). How *Candida* spp. evade neutrophil RNS is a major knowledge gap, critical to understand the pathogenesis of human disease. However, to overcome challenges associated with studying neutrophils *in vitro*, we require *in vivo* animal models to understand the neutrophil response in intact tissues.

Zebrafish are a highly tractable *in vivo* model system to investigate the behaviours of innate immune cells during infection (Henry *et al*., 2013; Speirs *et al*., 2024; Lam *et al*., 2004; Renshaw *et al*., 2006). Adaptive immunity does not mature until 4 to 6 weeks post fertilisation, whereas innate immune cells are present from early larval stages, allowing examination of innate immune responses to infections without the presence of adaptive immunity (Lam *et al*., 2004). Zebrafish models of *C. albicans* infection are well established, with a dose-dependent effect on zebrafish mortality (Chao *et al*., 2010), and have been used to uncover important disease mechanisms (Rosowski *et al*., 2018; Archambault *et al*., 2019; Blair *et al*., 2025). Furthermore, *C. albicans* is able to undergo the yeast-to-hyphae transition in zebrafish larvae, demonstrating a comparable pathogenesis to human infection (Chao *et al*., 2010; Gratacap *et al*., 2017; Seman *et al*., 2018; Christou *et al*., 2025).

Stimulation of RNS has the potential to be a host directed therapy to *C. albicans* infection that would complement existing antifungals. We have previously shown that Hypoxia inducible factor (Hif)-1α increases neutrophil RNS output, which is protective against the bacteria *Mycobacterium marinum* infection in zebrafish (Elks *et al*., 2013). HIF proteins are important regulators of cellular responses to hypoxia, innate immunity and inflammation (McGettrick and O’Neill, 2020). HIF-1α proteins are stabilised in hypoxic conditions, or by TLR signalling in normoxia, triggering transcription of target genes that regulate cellular metabolism, proliferation, migration, differentiation, angiogenesis and pro-inflammatory mediators (Peyssonnaux *et al*., 2005; Jantsch *et al*., 2011; Krzywinska and Stockmann, 2018; Hammond, Lewis and Elks, 2020). Therefore, we set out to address whether Hif-1α stabilisation may be a potential therapeutic mechanism that can enhance neutrophil RNS during *C. albicans* infections.

Here, we show that *C. albicans* infection suppressed neutrophil RNS in zebrafish larvae. Importantly, a similar dampening effect on RNS by *C. albicans* was also observed in human primary neutrophils. RNS suppression was not limited to laboratory strains of *C. albicans*; it was found during infection with *C. albicans* clinical isolates from non-invasive and invasive disease and with other *Candida* species. Heatkilled *C. albicans* was only partially able to decrease neutrophil RNS. Significantly, the ability of strains to reduce neutrophil RNS positively correlated with virulence *in vivo*. Both fungal arginase (car1) and host arginase induction were found to contribute to RNS reduction, with a *car1*Δ mutant having a reduced effect on neutrophil RNS and host arginase induced in zebrafish. Upregulation of host RNS production via Hif-1α stabilisation enhanced host control of *C. albicans* infection. Restoration of neutrophil RNS had an additive effect on host survival and clearance of *C. albicans* infection when combined with antifungals. Together, these data highlight RNS and Hif-1α as potential targets for host directed therapies against *C. albicans* infections.

## Methods

### Ethics Statement

Zebrafish were handled in accordance with Animals (Scientific Procedures) Act 1986, under Project Licenses P1A4A7A5E and PP7684817. Ethical approval was granted by the University of Sheffield Local Ethical Review Panel. Human blood was collected from healthy individuals following informed consent. Ethical approval was granted by the University of Sheffield Research Ethics Committee (study reference number 031773).

### Zebrafish Husbandry

Zebrafish were maintained in Home Office approved facilities in Biological Services Unit (BSU) aquaria at the University of Sheffield, in accordance with standard protocols and local animal welfare regulations. Adult fish were maintained at 28°C with a 14/10 hour light/dark cycle. Larvae were maintained at 28°C with a 14/10 hour light/dark cycle in 1X E3 medium (E3 60X stock: 5.0 mM NaCl, 0.17 mM KCl, 0.33 mM CaCl2, 0.33 mM MgSO4; diluted to 1X in distilled water) with 0.001% Methylene Blue (Sigma-Aldrich) to prevent fungal growth in the first 24 hours. From 24hpf, larvae were kept in E3 without Methylene Blue so that *Candida* spp. infection was not impacted. The following zebrafish strains were used: nacre (wild type), *Tg(mpx:GFP)i114* (Renshaw *et al*., 2006), *Tg(lyz:nfsB.mCherry)sh260* (Buchan *et al*., 2019), *Tg(mpeg:nlsClover*)sh436 (Bernut *et al*., 2019), *TgBAC(arg2:GFP)sh571* (Hammond *et al*., 2023).

### *Candida* spp. Culture

*C. albicans* strains (Table 1) and *Candida* spp. clinical isolates (Table 2) were used in this study. *C. albicans* was initially grown on YPD (4001022, MP Bio) plates at 28°C for 48 hours, then transferred to overnight liquid culture in YPD broth in a shaking incubator at 30°C at 200 rpm. *C. albicans* was heat-killed by incubation in a heat block at 65°C for 1 hour (Panpetch *et al*., 2017). To denature fungal proteins, *C. albicans* was boiled at 100°C for 5 minutes.

**Table 1:**
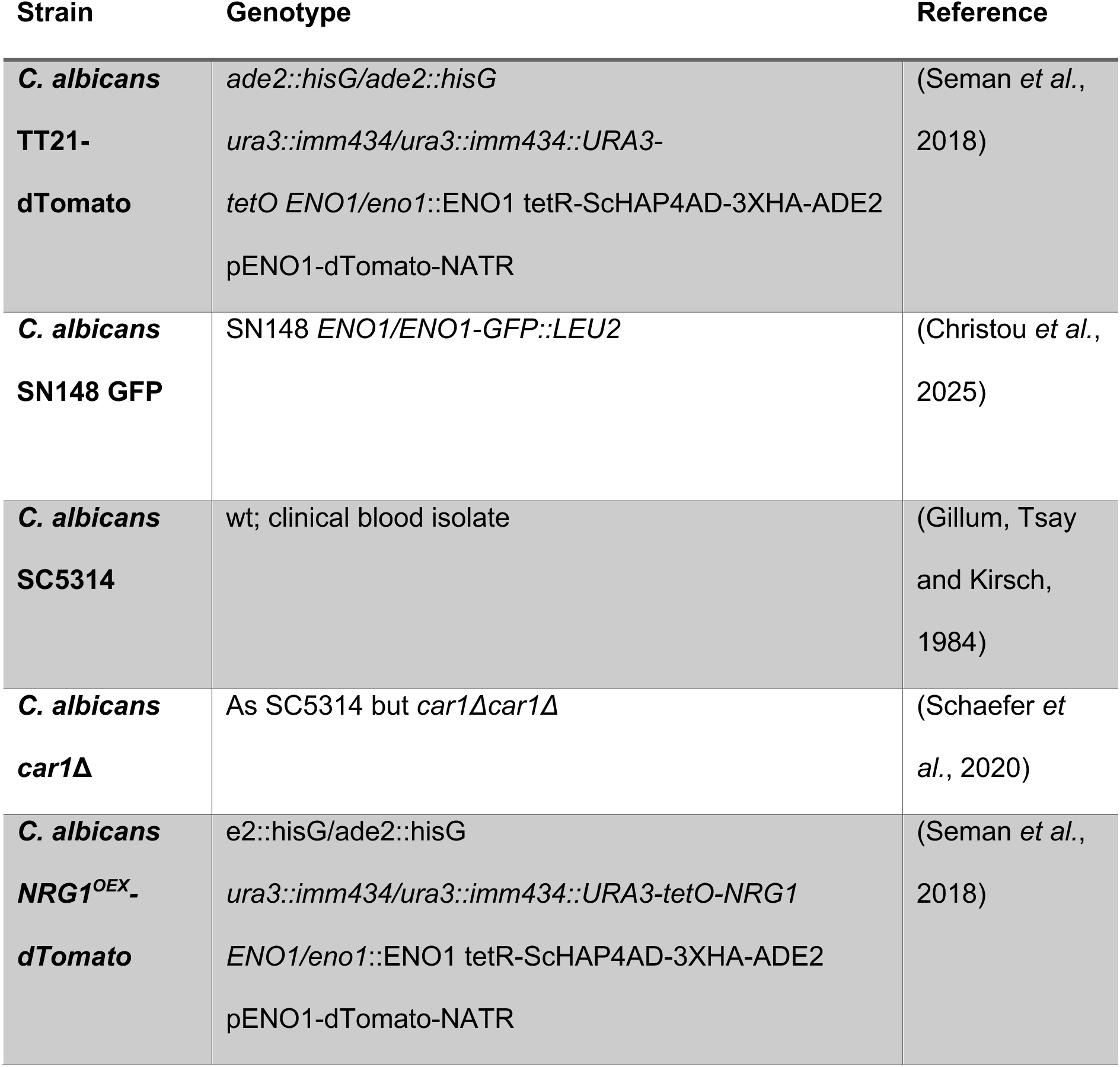
List of *C. albicans* strains

**Table 2:** List of *Candida* spp. clinical isolates

| Species | Name | Invasive/non-invasive | Sample Site | Source |
| --- | --- | --- | --- | --- |
| *Candida albicans* | AJP4 | Non-invasive | Vaginal sample | Provided by Dr David Partridge |
| *Candida albicans* | AJP5 | Invasive | Blood sample | Collected at Sheffield Children's Hospital |
| *Candida albicans* | AJP9 | Non-invasive | Tongue swab | Collected at Sheffield Children's Hospital |
| *Candida albicans* | AJP25 | Invasive | Line tip sample | Collected at Sheffield Children's Hospital |
| *Candida glabrata* | AJP12 | Invasive | Urine sample | Collected at Sheffield Children's Hospital |
| *Candida parapsilosis* | AJP22 | Invasive | Urine sample | Collected at Sheffield Children's Hospital |
| *Candida guilliermondii* | AJP24 | Non-invasive | Tracheal swab | Collected at Sheffield Children's Hospital |
| *Candida auris* | StG2 | Invasive | Bloodstream | Provided by Dr Tihana Bicanic |

### C. albicans Infections

Overnight liquid *C. albicans* cultures were washed 3 times in phosphate buffered saline (PBS) and counted on a haemocytometer. Washed cultures were resuspended in 10% PVP (polyvinylpyrrolidone; Calbiochem) to achieve the desired inoculation. Infection dose for systemic caudal vein infections were 200 colony forming units (cfu) in a 1nl injection volume, 100 cfu in 1nl for experiments with immune cell ablation or 500 cfu in 1nl in experiments with antifungals.

For survival curves, zebrafish survival was monitored daily and dead zebrafish were removed. Death was determined by a cessation of heartbeat/circulation or tissue degradation post-mortem. At 4 days post infection (dpi; equivalent to 5 days post fertilisation, dpf), all surviving zebrafish larvae were culled.

### RNA Injections

Larvae were injected with dominant active *hif-1αb* variant RNA (DA1) or dominant negative *hif-1αb* variant RNA in dH2O with 10% phenol red (Sigma-Aldrich, to visualise successful injection) at the one-cell stage, as previously described (Elks *et al*., 2011). 10% (v/v) phenol red in dH2O (PR; Sigma-Aldrich) was used as a solvent control.

### Morpholino Injections

A *csf3r* morpholino (Gene Tools) was used to deplete neutrophil populations (Ellett *et al*., 2011). A standard control morpholino (Gene Tools) used as a negative control. The antisense morpholino oligonucleotide sequences were: *csf3r* – 5’-GAAGCACAAGCGAGACGGATGCCAT-3’; control – 5’-CCTCTTACCTCAGTTACAATTTATA-3’. *Csf3r* morpholino was functionally validated by whole body neutrophil count at 2 dpf.

### F0 CRISPR-Cas9 Knockdown

Two sgRNAs were designed to target the ATG/first exon of *nos2a* and *nos2b* to knockdown zebrafish *nos2* genes by CRISPR-Cas9. Efficacy of knockdown of RNS production was functionally validated by whole-body anti-nitrotyrosine staining. Tyrosinase (*tyr*) sgRNA, targeting a pigment gene that plays no role in neutrophil RNS production, was used as a negative control and to demonstrate Cas9 function (Isles *et al*., 2019). The target sequences were: *nos2a* – 5’-TTTCTCATTTTCAATGATAG-3’; *nos2b* – 5’-GTTCGCTCTTGTGAGTGACC-3’; *tyr* – 5’-CCTGACCTCCTGAAGACCCC-3’. 25μM sgRNA was co-injected with tracrRNA, 1 μl Cas9 and phenol red.

### Pharmacological Treatment of Zebrafish Larvae

For pharmacological inhibition of nitric oxide synthase, larvae were treated with 200 μM N6-(1-Iminoethyl)-lysine (L-NIL) by immersion. dH2O was used as a solvent control.

For antifungal survival curves and clearance assays, larvae were treated with either 5.0 µg/ml fluconazole in DMSO or 0.5 µg/ml caspofungin in dH2O by immersion. DMSO or dH2O was used as a solvent control. At 2 dpi, larvae were transferred into fresh E3 and treatment was re-administered.

### Anti-nitrotyrosine Staining

1 dpi larvae (2 dpf) were fixed in 4% formaldehyde in PBS overnight at 4°C. Whole-body nitrotyrosine levels were labelled using a rabbit polyclonal anti-nitrotyrosine antibody (06-284; Merck Millipore) and were detected using Alexa-633 conjugated goat-anti-rabbit secondary antibody (Invitrogen Life Technologies), as previously described (Elks *et al*., 2013, 2014) in a T*g(mpx:GFP)i114* background to assess for neutrophil specific RNS.

### Microscopy and Quantification of Anti-nitrotyrosine Staining

For *C. albicans* SC5314 and *car1*Δ, brightfield images were taken using an inverted Leica DMi8 with a 40x lens and a Hamamatsu OrcaV4 camera. Stained larvae were imaged on a Leica DMi8 inverted microscope with a Leica TCS-SPE line-scanning confocal for imaging with a 40x 1.1NA water immersion lens. For quantification purposes, acquisition settings were kept the same across the groups. Corrected fluorescence intensity was calculated using FIJI measurements assessing the cell fluorescence of individual cells corrected for cell size and background fluorescence of the image (Burgess *et al*., 2010; Elks *et al*., 2013, 2014).

### Hyphal staging

For assessment of fungal growth of *Candida* spp clinical isolates, which lack fluorescent proteins, brightfield images were taken using an inverted Leica DMi8 with a 40x lens and a Hamamatsu OrcaV4 camera. Classification of hyphal stages was done manually, based on a previously described descriptive criteria (Thrikawala *et al*., 2022). Criteria for hyphal staging were:

1. Yeast: only spherical, yeast cells were observed. No signs of any hyphal growth.
2. Germinating: ovoid cells, resembling pseudohyphae or very short hyphae, were observed.
3. Hyphal extension: a small number of short or medium length hyphae were observed.
4. Destructive hyphae: multiple invasive hyphae were observed. Majority of observed *C. albicans* was in hyphal morphotype.

### Human neutrophils

Human neutrophils were isolated by Plasma-Percoll density gradient centrifugation of whole blood from healthy donors as previously described (Herman, Rahman and Prince, 2020). 0.5x 10^6^ cells were added to individual wells in 96-well plates in 100μl volumes. Cells were primed with 1 μg/ml lipopolysaccharides from *Escherichia coli* (LPS—Sigma, L8274). Neutrophils were infected with an MOI of 1.0 or 0.5 *Candida albicans*, MOI 1.0 heat-killed *Candida albicans* or uninfected (sterile RPMI media) and were incubated at 37°C, 5% CO2 for 3 hours. Cells were cytocentrifuged onto microscope slides and fixed in 4% formaldehyde for staining. Immunostaining was performed as described above.

### Microscopy of *C. albicans* Infection clearance

Brightfield images were taken using an inverted Leica DMi8 with a 2.5x lens and a Hamamatsu OrcaV4 camera. Surviving fish were classified as ‘infection cleared’ only if no signs of remaining fluorescent *C. albicans* infection could be observed at 2.5x magnification.

### Statistical Analysis

All data were analysed using GraphPad Prism 10.5.0 1 (GraphPad Software, La Jolla, California, USA, www.graphpad.com). Nitrotyrosine staining data was analysed with two-tailed Mann-Whitney test or Kruskal-Wallis test, with Dunn’s multiple comparisons test. Survival curves were analysed with Gehan-Breslow-Wilcoxon test, with Bonferroni correction. The p values shown are: ns=not significant, *p<0.05, **p<0.01, ***p<0.001, ****p<0.0001

## Results

### *C. albicans* infection suppresses neutrophil RNS production below basal levels

Live *C. albicans* caused a 90.0% reduction in the levels of RNS (visualised using an anti-nitrotyrosine antibody) compared to mock infection (PVP) controls (Figure 1A-B), indicating not only a lack of induction of RNS, but also a downregulation of basal RNS. We have previously shown that RNS is predominantly found in neutrophils both in unchallenged and mock infected (PVP) individuals (Elks *et al*., 2014) and is upregulated after bacterial challenge with *Mycobacterium marinum* (Elks *et al*., 2013). This established that *C. albicans* supressed neutrophil RNS *in vivo* in contrast to bacterial challenge. In mock infected (PVP) controls there was a basal level of nitrotyrosine levels in neutrophils, as observed previously (Elks *et al*., 2013, 2014). We tested another bacterium, *Staphylococcus aureus* to determine if induction of neutrophil RNS was limited to mycobacteria. *S. aureus* (SH1000) infection also caused a significant increase in neutrophil nitrotyrosine fluorescence, corroborating previous observations of increased RNS levels in response to mycobacterial infection (Figure S1; Elks *et al*., 2013). RNS reduction by *C. albicans* was not limited to a single strain as we infected with a different *C. albicans* strain, SN148 GFP, and observed similar RNS suppression (Figure S2). This allowed the use of TT21-dTomato and SN148 GFP interchangeably, depending on the colour combinations required for each experiment. Zebrafish larvae infected with heat-killed C. albicans TT21-dTomato had an intermediate level of RNS suppression, between the PVP control and live C. albicans infection (Figure 1A-B). Hence, the full effect of neutrophil RNS suppression is dependent on live C. albicans suggestive of partially active and passive mechanisms.

**Figure 1:**
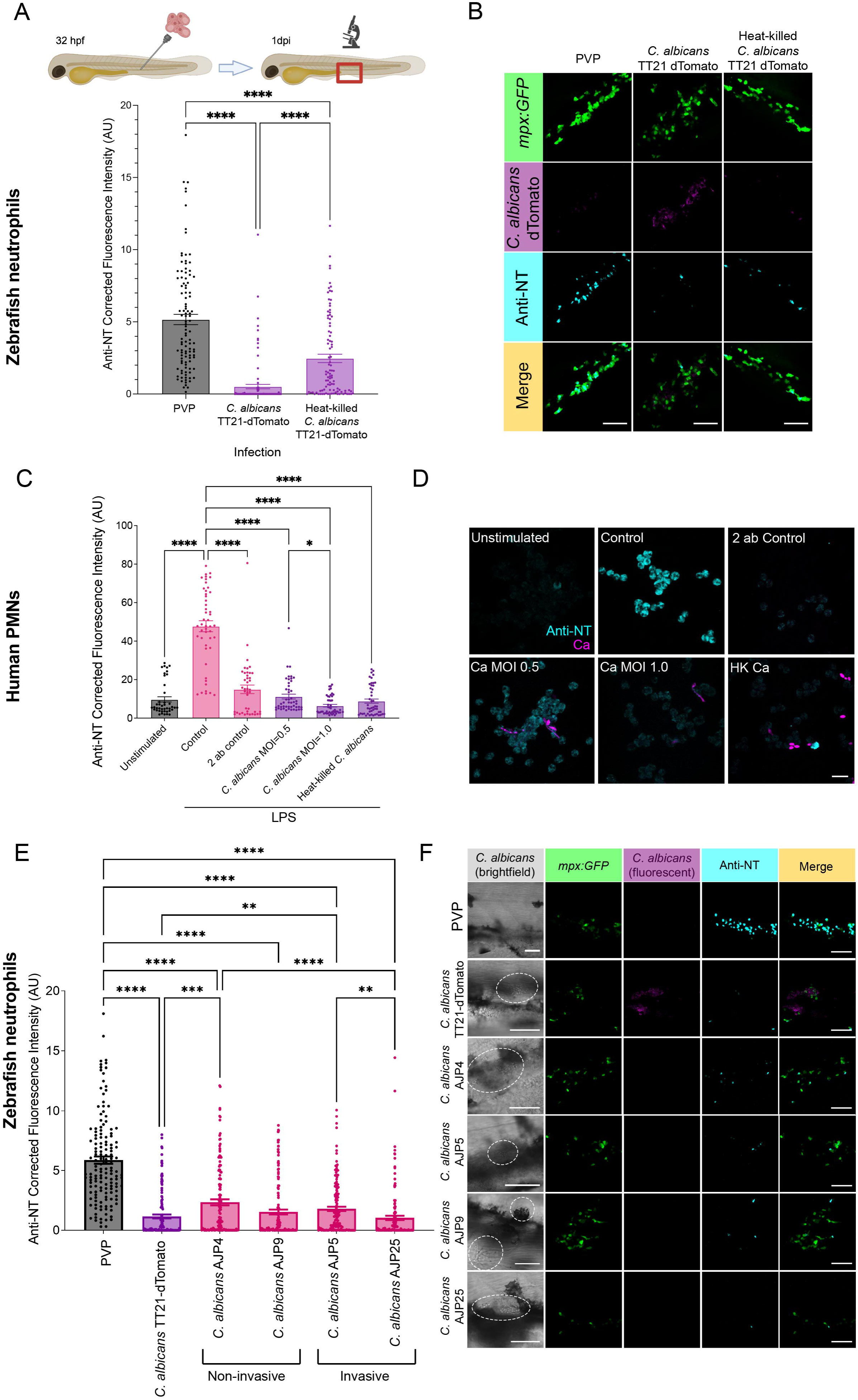
*Candida albicans* suppresses neutrophil reactive nitrogen species (RNS) production below resting levels. (A) Schematic of experiment: 1 dpf zebrafish were injected with PVP or *C. albicans* into the caudal vein. At 1 dpi, larvae were fixed and stained with anti-nitrotyrosine primary antibody and goat-anti-rabbit Alexa-633 secondary antibody. Zebrafish were then imaged at the site of infection in the caudal haematopoietic tissue (CHT) and anti-nitrotyrosine fluorescence was quantified. Graph shows anti-nitrotyrosine fluorescence at 1 dpi following injection of PVP, live *C. albicans* TT21-dTomato or heat-killed *C. albicans* TT21-dTomato into the caudal vein at 30 hpf. N=96-108 neutrophils from 16-18 fish, obtained from 3 independent experiments. Error bars show SEM. Statistical significance determined by Kruskal-Wallis test, then Dunn’s multiple comparisons test. P values shown: ****p<0.0001. (B) Representative images of PVP, *C. albicans* TT21-dTomato and heat-killed *C. albicans* TT21-dTomato infected zebrafish larvae at 1 dpi. Scale bars = 50 μm. (C) Corrected fluorescence intensity of PMNs fixed at 3 hours post LPS treatment and stained with anti-nitrotyrosine antibody (in arbitrary units, AU). Non-LPS stimulated resting neutrophils, LPS stimulated neutrophils and a secondary antibody only groups were included as controls. Neutrophils infected with an MOI of 0.5 and 1.0 live *C. albicans* and heat-killed *C. albicans* all show a decrease in nitrotyrosine in the presence of LPS compared to LPS-only controls. N=42-48 neutrophils from 2 independent experiments and donors. (D) Representative images of human PMNs stained with anti-NT. Scale bars = 50 μm. *p<0.05, ****p<0.0001. (E) Anti-nitrotyrosine fluorescence at 1 dpi following injection of PVP, *C. albicans* TT21-dTomato or *C. albicans* clinical isolates into the caudal vein at 30 hpf. N=144 neutrophils from 24 fish, obtained from 3 independent experiments. Error bars show SEM. Statistical significance determined by Kruskal-Wallis test, then Dunn’s multiple comparisons test. P values shown: **p<0.01, ***p<0.001, ****p<0.0001. (F) Representative images of PVP, *C. albicans* TT21-dTomato, *C. albicans* AJP4, *C. albicans* AJP5, *C. albicans* AJP9 and *C. albicans* AJP25 infected zebrafish larvae at 1 dpi. White dotted circles show *C. albicans.* Scale bars = 50 μm.

To confirm our finding in humans, we performed anti-nitrotyrosine staining on human primary polymorphonuclear neutrophils (PMNs) after *C. albicans* infection *in vitro*. Detectable and robust RNS induction in human neutrophils was induced by priming with LPS which led to a robust nitrotyrosine response after 3 hours in the absence of infection (PVP mock infected) (Figure 1C-D). When *C. albicans* was added at an MOI of 1.0 nitrotyrosine was suppressed to baseline unstimulated levels, despite the presence of LPS (Figure 1C-D). A lower MOI of 0.5 live *C. albicans* showed less of an RNS suppression compared to an MOI of 1.0 of live *C. albicans*, indicating a dose dependent effect (Figure 1C-D). Similarly, an MOI of 1.0 of heat-killed *C. albicans* exhibited lower RNS suppression compared to the same MOI of live *C. albicans*. Together these data show that *C. albicans* suppresses RNS production in human neutrophils as in zebrafish.

The laboratory reference strain SC5314 and strains derived from this (including SN148 and TT21) have inactivating missense mutations in RNAi component Argonaute and so may behave differently to *C. albicans* strains found in the clinic (Iracane *et al*., 2024). Therefore, clinical isolates from invasive (AJP5 and AJP25) and non-invasive (AJP4 and AJP9) infections were tested for neutrophil RNS suppression in zebrafish. Our clinical isolates lacked a fluorescent protein marker, so brightfield microscopy was used to identify sites of infection in anti-nitrotyrosine stained zebrafish larvae. Both invasive (AJP5 and AJP25) and non-invasive (AJP4 and AJP9) isolates were able to significantly reduce neutrophil RNS compared to mock infection (PVP) controls (Figure 1E-F). These observations demonstrate neutrophil RNS suppression by *C. albicans* is conserved in clinical isolates, implying RNS suppression in *C. albicans* infections may be a clinically relevant phenomenon.

### Full neutrophil RNS suppression is dependent on hypha-forming *C. albicans*

*C. albicans* morphological switch from yeast to hyphae is associated with increased virulence and shifts in the transcriptome and secretome (Nantel *et al*., 2002; Höfs, Mogavero and Hube, 2016; Vaz *et al*., 2021). Classification of hyphal growth by *C. albicans* TT21 and clinical isolates in zebrafish revealed the isolates that caused greater neutrophil RNS suppression (AJP9 and AJP25) (Figure 1E-F) exhibited greater hyphal growth *in vivo* compared to those that reduced RNS to a lesser extent (AJP4 and AJP5) (Figure 2A-B). Based on this, we hypothesised hyphal switching may be partially responsible for the suppression of neutrophil RNS. To investigate this, 1 dpf zebrafish were infected with *C. albicans* TT21-dTomato or yeast-locked strain *C. albicans* NRG1^OEX^-dTomato (Seman *et al*., 2018). Both *C*. *albicans* NRG1^OEX^-dTomato and heat-killed *C. albicans* NRG1^OEX^-dTomato caused a modest decrease in RNS compared to wildtype strains, with reductions comparable to heat-killed *C. albicans* (Figure 2C-D). These data suggest that an important component of full RNS suppression by *C. albicans* is hyphal switch-dependent, but that hyphal switch is not essential for neutrophil RNS suppression.

**Figure 2:**
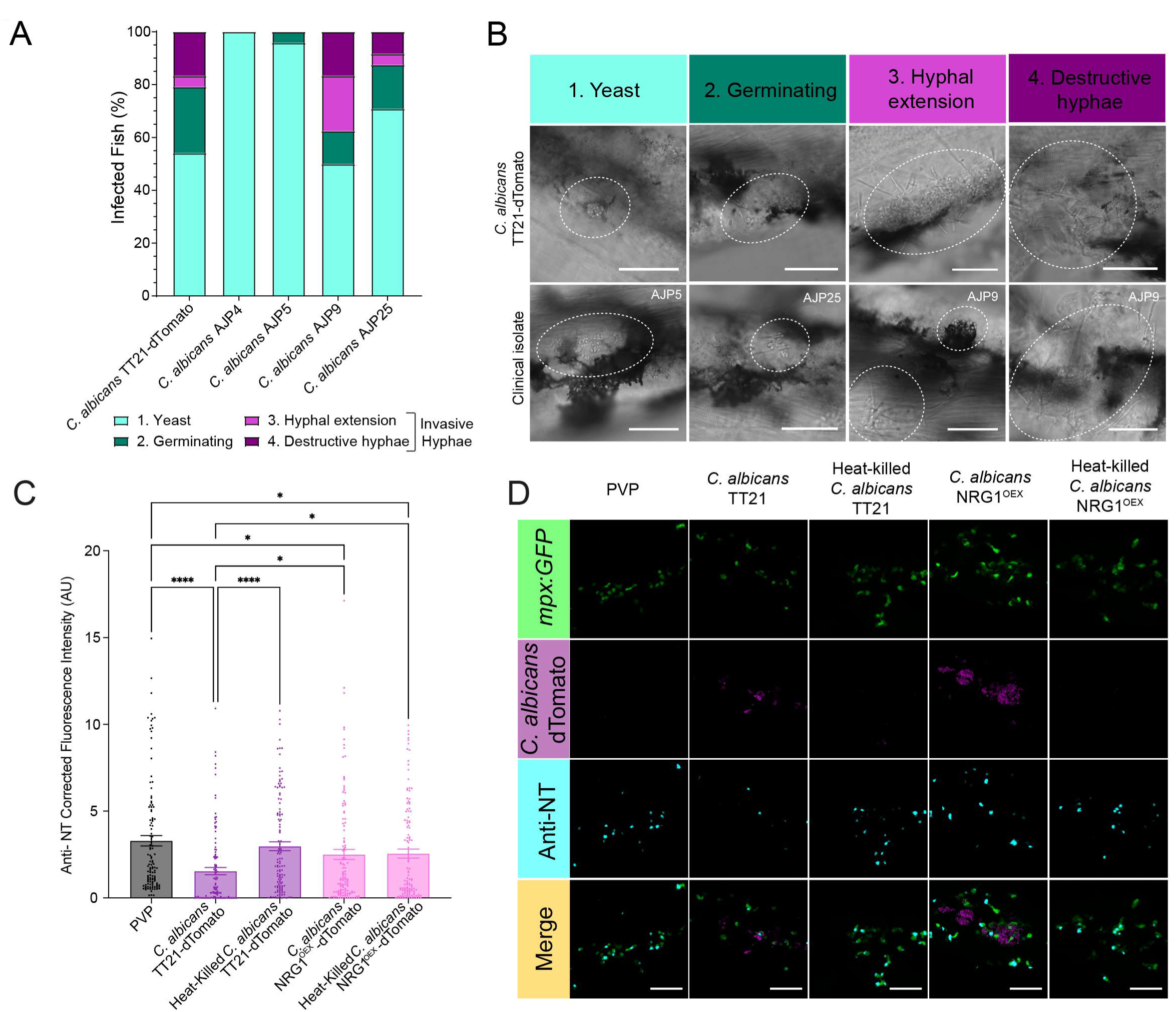
Robust RNS suppression is associated with hyphae-forming *C. albicans* (A) Hyphal classification of zebrafish larvae infected with range of *C. albicans* isolates at 1 dpi. N=24 fish, obtained from 3 independent experiments. (B) Representative images of hyphal classification in 1 dpi zebrafish larvae infected with *C. albicans* TT21-dTomato or *C. albicans* clinical isolate. Images are labelled with the *C. albicans* isolate. White dotted circles show regions of fungal growth. Scale bar = 50 μm. (C) Anti-nitrotyrosine fluorescence at 1 dpi following injection of PVP, *C. albicans* TT21-dTomato, heat-killed *C. albicans* TT21-dTomato*, C. albicans* NRG1^OEX^-dTomato or heat-killed *C. albicans* NRG1^OEX^-dTomato into the caudal vein. N=120 neutrophils from 20 fish, obtained from 3 independent experiments. Error bars show SEM. Statistical significance determined by Kruskal-Wallis test, with Dunn’s multiple comparisons tests. *p<0.05, ****p<0.0001. (D) Representative images of PVP, *C. albicans* TT21-dTomato, heat-killed *C. albicans* TT21-dTomato*, C. albicans* NRG1^OEX^-dTomato or heat-killed *C. albicans* NRG1^OEX^-dTomato infected larvae at 1 dpi. Scale bars = 50 μm.

### Host arginase is induced by *C. albicans*

Inducible nitric oxide synthase (iNOS; Nos2 in zebrafish) is the enzyme responsible for RNS production. iNOS competes with the arginase enzyme for a shared substrate, L-arginine, and arginase can competitively inhibit RNS production by iNOS (Bansal and Ochoa, 2003). We investigated the impact of *C. albicans* infection on host arginase levels, as a potential mechanism by which neutrophil RNS is reduced. PVP injected larvae had a low basal level of *arg2:GFP* expression (Figure 3A-B). *Mycobacterium marinum* (used as a control for arginase induction) caused a significant upregulation of *arg2:GFP* compared to PVP, as previously described (Hammond et al., 2023). *C. albicans* TT21-dTomato caused a greater upregulation in *arg2:GFP* expression compared to *M. marinum* indicating a robust host arginase upregulation. *C. albicans* NRG1^OEX^-dTomato also caused upregulation of *arg2:GFP* compared to both PVP and *M. marinum* indicating this is a hyphal-independent process. Examination of *arg2:GFP* expression in neutrophils (labelled by *mpx:GFP*) revealed *C. albicans* TT21-dTomato caused an increase in the percentage of *arg2:GFP+* neutrophils (41.59% of the neutrophil population in field of view; Figure 4C-D) compared to PVP (2.01%) and *M. marinum* (10.97%). Together, these results reveal *C. albicans* infection upregulates host arginase expression in a subpopulation of neutrophils to a greater level than the professional bacterial pathogen *M. marinum*.

**Figure 3:**
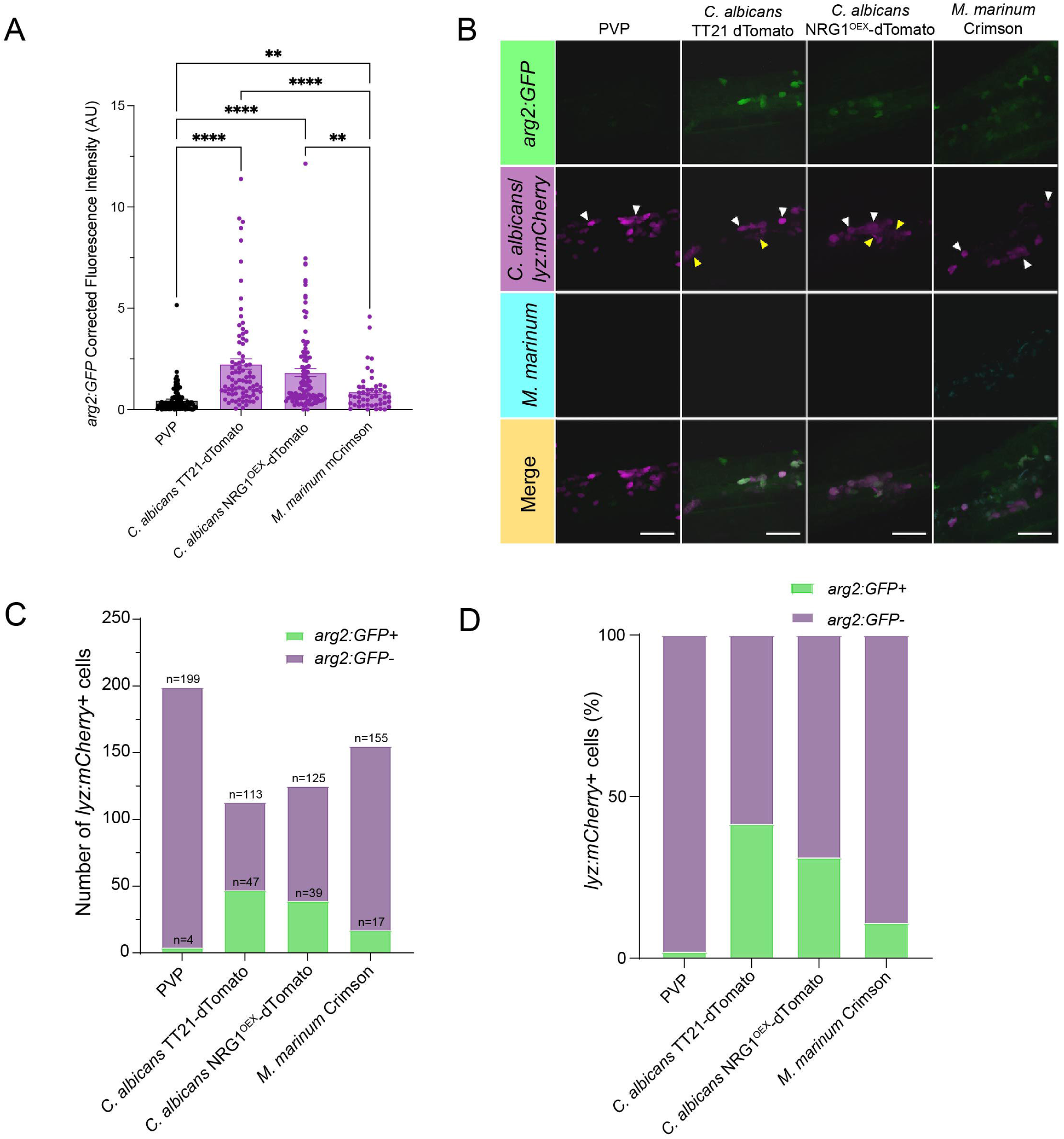
*C. albicans* infection stimulates host *arginase 2* expression. (A) Corrected fluorescence intensity of *Tg(arg2:GFP)* zebrafish larvae at 1 dpi following injection of PVP, *C. albicans* TT21-dTomato, *C. albicans* NRG1^OEX^-dTomato or *M. marinum* Crimson into the caudal vein at 30 hpf. For PVP, *C. albicans* TT21-dTomato and *C. albicans* NRG1^OEX^-dTomato: n=84-108 cells from 14-18 fish, obtained from 3 independent experiments. For *M. marinum*: n=48 cells from 8 fish, obtained from 2 independent experiments. Error bars show SEM. Statistical significance determined by Kruskal-Wallis test, then Dunn’s multiple comparisons test. P values shown: **p<0.01, ****p<0.0001 (B) Representative images of *Tg(arg2:GFP)* zebrafish embryos injected with PVP, *C. albicans* TT21-dTomato, *C. albicans* NRG1^OEX^-dTomato or *M. marinum* Crimson at 24hpi. White arrow heads point towards neutrophils. Yellow arrow heads point towards *C. albicans*. Scale bars = 50 μm. (C) Number of *lyz:nfsB.mCherry*-expressing cells that are *arg2:GFP*+ and *arg2:GFP-,* following infection with *C. albicans* TT21-dTomato, *C. albicans* NRG1^OEX^-dTomato, *M. marinum* Crimson or PVP. N=113-199 cells from 12 zebrafish, obtained from 2 independent experiments. (D) Percentage of *lyz:nfsB.mCherry*-expressing neutrophils that are *arg2:GFP*+ and *arg2:GFP-,* following infection with *C. albicans* TT21-dTomato, *C. albicans* NRG1^OEX^-dTomato, *M. marinum* Crimson or PVP. N=113-199 cells from 12 zebrafish, obtained from 2 independent experiments.

**Figure 4:**
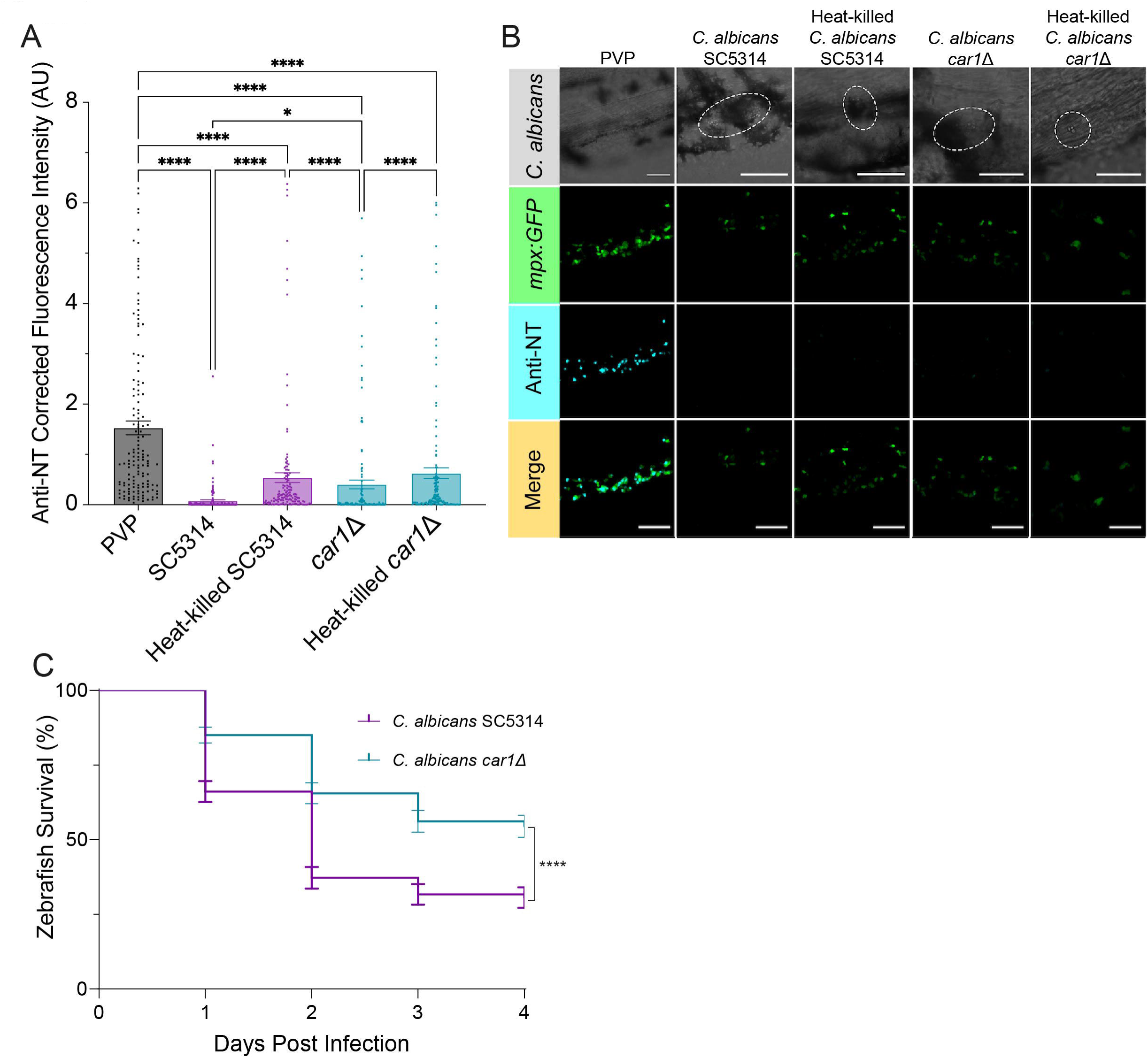
*C. albicans car1* contributes to suppression of neutrophil RNS. (A) Anti-nitrotyrosine fluorescence at 1 dpi following injection of PVP, *C. albicans* SC5314, heat-killed *C. albicans* SC5314*, C. albicans car1*Δ or heat-killed *C. albicans car1*Δ into the caudal vein. N=144 neutrophils from 24 fish, obtained from 3 independent experiments. Error bars show SEM. Statistical significance determined by Kruskal-Wallis test, with Dunn’s multiple comparisons tests. *p<0.05, ****p<0.0001. (B) Representative images of anti-nitrotyrosine stained larvae. Dashed lines show *C. albicans*. Scale bars = 50 μm. (C) 1 dpf nacre larvae were injected into the caudal vein with 200 cfu *C. albicans* SC5314 or *C. albicans car1Δ*. Larval survival was measured daily up to 4 dpi. n=180 fish, obtained from 3 independent experiments. Error bars show SEM. Statistical significance determined by Gehan-Breslow-Wilcoxon test, with Bonferroni correction. P values shown: ****p<0.0001

### *C. albicans* derived arginase contributes to the suppression of neutrophil RNS

Fungi are known to produce enzymes with similar structure and functions to vertebrate enzymes. *C. albicans* produces its own version of arginase, *CAR1* (Schaefer *et al*., 2020). We hypothesised that *C. albicans* arginase interfered with the host RNS production, resulting in neutrophil RNS suppression. 1 dpf zebrafish were infected with *C. albicans car1*Δ or its parental strain *C. albicans* SC5314 and RNS production was measured at 1 dpi by anti-nitrotyrosine staining. *C. albicans car1*Δ infection decreased neutrophil RNS production compared to PVP (Figure 4A-B). *C. albicans car1*Δ caused an intermediate level of RNS suppression between live *C. albicans* SC5314 and heat-killed *C. albicans* SC5314 (Figure 4A-B), indicating that CAR1 contributes to the suppression of neutrophil RNS. Heat-killed *C. albicans car1*Δ was not significantly different from heat-killed *C. albicans* SC5314. Infection with *C. albicans car1Δ* resulted in significantly greater zebrafish larvae survival than infection with *C. albicans* SC5314 (Figure 4C), suggesting RNS suppression by car1 is important for *C. albicans* virulence *in vivo*.

### The level of neutrophil RNS suppression by *Candida* spp. correlates with virulence

*C. albicans* is the leading causative agent of candidiasis, but there are many other clinically important *Candida* spp. We investigated the effect of a range of *Candida* spp. clinical isolates on neutrophil RNS levels *in vivo*. Clinical isolates of *C. parapsilosis* (AJP22), *C. guilliermondii* (AJP24) and *C. auris* (StG2) all robustly reduced neutrophil RNS compared to PVP controls, but to a lesser extent than a laboratory strain of *C. albicans* (Figure 5A-B). *C. glabrata* AJP12 caused a more modest decrease in neutrophil RNS compared to *C. albicans*. (Figure 5A-B). Together, these results show that neutrophil RNS suppression is conserved across *Candida* spp. clinical isolates, but that levels of RNS suppression can vary between species.

**Figure 5:**
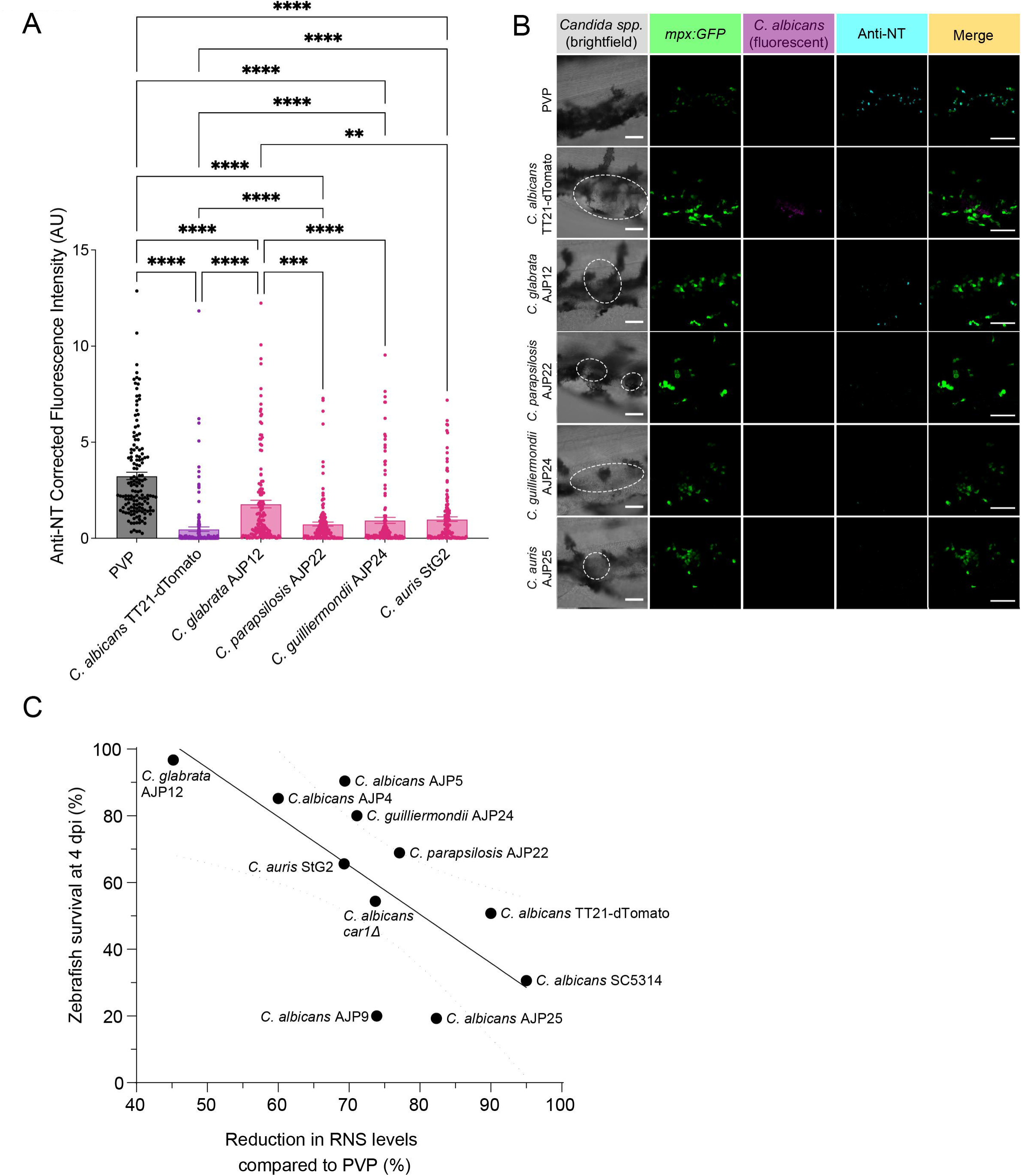
Non-albicans *Candida* spp. suppress neutrophil RNS. (A) Anti-nitrotyrosine fluorescence at 1 dpi following injection of PVP, *C. albicans* TT21-dTomato, *C. glabrata* AJP12, *C. parapsilosis* AJP22, *C. guilliermondii* AJP24 or *C. auris* StG2 into the caudal vein. N=144 neutrophils from 24 fish, obtained from 3 independent experiments. Error bars show SEM. Statistical significance determined by Kruskal-Wallis test, with Dunn’s multiple comparisons tests. * indicates difference compared to PVP. † indicates difference compared to *C. albicans* TT21-dTomato. ‡ indicates difference compared to *C. glabrata* AJP12. P values shown: **p<0.01, ***p<0.001, ****p<0.0001. (B) Representative images of PVP, *C. albicans* TT21-dTomato, *C. glabrata* AJP12, *C. parapsilosis* AJP22, *C. guilliermondii* AJP24 or *C. auris* StG2 infected larvae at 1 dpi. Scale bars = 50 μm. (C) Reduction in RNS levels compared to PVP control plotted against zebrafish survival at 4dpi, following systemic infection with 200 cfu *Candida* spp. Full survival curves for *Candida* spp. are in Figures S4 and S5. Each plotted point is based on 3 independent repeats of relevant experiment. Line of best fit and 95% confidence error lines are shown. Pearson’s correlation coefficient and R^2^ value were calculated.

When percentage zebrafish survival at 4 dpi were plotted against the percentage reduction of RNS levels by *C. albicans* strains/isolates compared to PVP control, survival negatively correlated with reduction in neutrophil RNS levels (Figure 5C. Full survival curves for *Candida* spp. are in Figure S3 and S4.) (Pearson’s correlation coefficient: -0.718, 95% confidence interval: -0.208 – -0.921; p<0.05, R^2^ value: 0.516). Therefore, a greater suppression of host RNS levels is correlated with increased virulence in zebrafish infection *in vivo*, indicating that suppression of neutrophil RNS is a potential virulence mechanism in human disease-relevant *Candida* spp..

### Increased neutrophil RNS, via Hif-1α stabilisation, is protective against *C. albicans* infection

Hif-1α is a transcription factor with known roles in regulation of innate immunity (McGettrick and O’Neill, 2020). We have previously shown stabilised Hif-1α (using injection of a dominant active variant RNA) is protective against *M. marinum* infection *in vivo* by increasing neutrophil RNS levels (Elks *et al*., 2013; Hammond, Lewis and Elks, 2020). We therefore hypothesised that stabilisation of Hif-1α would restore neutrophil RNS in *C. albicans* infection and increase host survival.

1 cell stage zebrafish larvae were injected with water/10 % phenol red (solvent) control (PR), dominant negative *hif-1α* (DN *hif-1α*) or dominant active *hif-1α* (DA *hif-1α*) RNA (as previously described; (Elks *et al*., 2011)), and subsequently infected systemically with *C. albicans* TT21-dTomato infection at 30 hpf. DA *hif-1α* larvae had significantly greater survival over the first 4 days post infection than both PR and DN *hif-1α* larvae (Figure 6A), demonstrating that Hif-1α stabilisation is host-protective in *C. albicans* infection.

**Figure 6:**
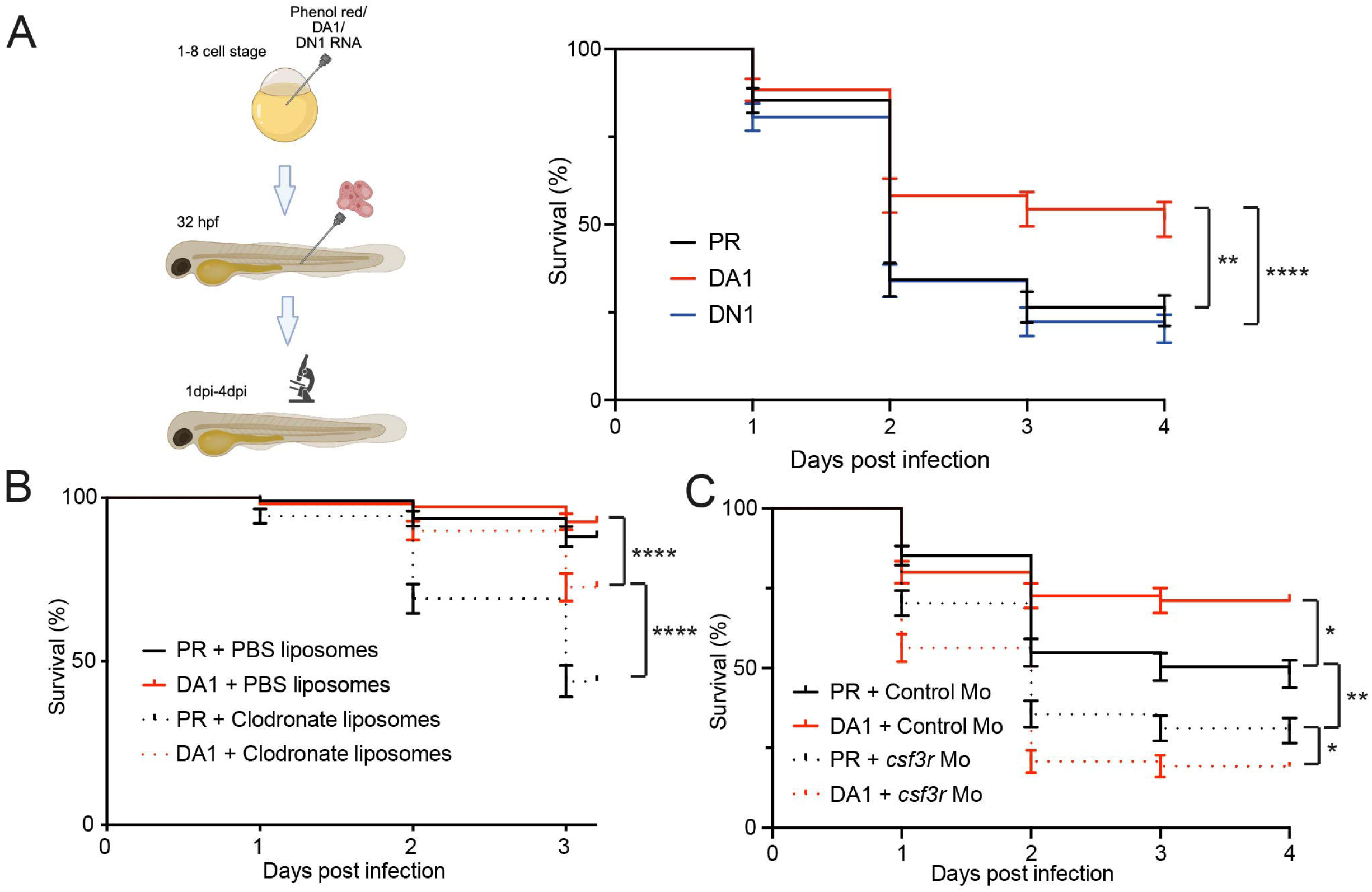
Hif-1α stabilisation is protective against *C. albicans* infection. (A) Survival of *C. albicans* TT21-dTomato infected zebrafish larvae following injection with dominant active *hif-1α* (DA1), dominant negative *hif-1α* (DN1) RNA or phenol red (PR) control. Mortality was measured daily. N=102-103 fish, obtained from 3 independent experiments. Error bars show SEM. Statistical significance determined by Gehan-Breslow-Wilcoxon test with Bonferroni correction. **p<0.01, ****p<0.0001. (B) Survival curve of 100 cfu *C. albicans* TT21-dTomato infected zebrafish larvae following injection with DA1 or PR, then treatment with PBS or clodronate liposomes. Mortality was measured daily. N=102-103 fish, obtained from 3 independent experiments. Error bars show SEM. Statistical significance determined by Gehan-Breslow-Wilcoxon test with Bonferroni correction. ****p<0.0001. (C) Survival curve of 100 cfu *C. albicans* TT21-dTomato infected zebrafish larvae following injection with PR or DA1 and control or *csf3r* morpholino. Mortality was measured daily. N=135 fish, obtained from 3 independent experiments. Error bars show SEM. Statistical significance determined by Gehan-Breslow-Wilcoxon test with Bonferroni correction. *p<0.05, **p<0.01.

Neutrophils play an essential role in innate immune control of invasive candidiasis (Desai and Lionakis, 2018). In zebrafish neutrophils are responsible for the killing of *C. albicans*, whereas macrophages internalise but do not kill (Brothers *et al*., 2013). Therefore, we hypothesised that the protective effect of Hif-1α stabilisation would be mediated by neutrophils rather than macrophages. Firstly, to assess the impact of Hif-1α stabilisation on macrophage-mediated immunity, we depleted the macrophage population by injection of clodronate liposomes (van Rooijen and Hendrikx, 2010) at 30 hpf (or PBS liposomes as a negative control) leading to a significant depletion of the macrophage population (Figure S5). At 30 hpf, PR or DA *hif-1α* larvae were injected into the caudal vein with clodronate liposomes or PBS liposomes followed by infection with 100cfu *C. albicans* TT21-dTomato at 2 dpf. Both groups of PBS liposome-injected larvae had greater survival than their clodronate liposome, macrophage-depleted, siblings. DA *hif-1α* was host protective both with or without depletion of macrophages, with DA *hif-1α* + clodronate liposome larvae having greater survival than PR + clodronate liposome larvae (Figure 6B; p<0.0001), indicating that the host-protective effect of Hif-1α stabilisation is not dependent on the presence of macrophages. Next, we injected 1 cell stage zebrafish larvae with a *csf3r* morpholino (Ellett *et al*., 2011) to deplete neutrophil numbers by over 60% in 2 dpf zebrafish larvae (Figure S6). *csf3r* morpholino injection led to a significant decrease in survival of 100 cfu *C. albicans* infected larvae in both PR and DA *hif-1α* larvae (Figure 6C). DA *hif-1α* + *csf3r* larvae had lower survival compared to PR + *csf3r* larvae, showing a loss of the protective effect of Hif-1α stabilisation in neutrophil ablated larvae (Figure 6C), indicating that the protective effect of Hif-1α stabilisation is neutrophil-dependent. DA *hif-1α* did not alter the recruitment of neutrophils to the site of a local hindbrain *C. albicans* infection compared to PR injected controls (Figure S7), therefore we hypothesised that the host protective effect of Hif-1α stabilisation in *C. albicans* infection was due to increased neutrophil RNS production. DA *hif-1α* doubled the levels of nitrotyrosine compared to PR in mock infected (PVP) larvae (Figure 7A-B) similar to previous observations (Elks *et al*., 2013). While *C. albicans* reduced nitrotyrosine levels in PR control injected larvae, DA *hif-1α C. albicans* infected larvae had high levels of neutrophil nitrotyrosine compared to mock infected (PVP) DA1 controls (Figure 7A-B).

**Figure 7:**
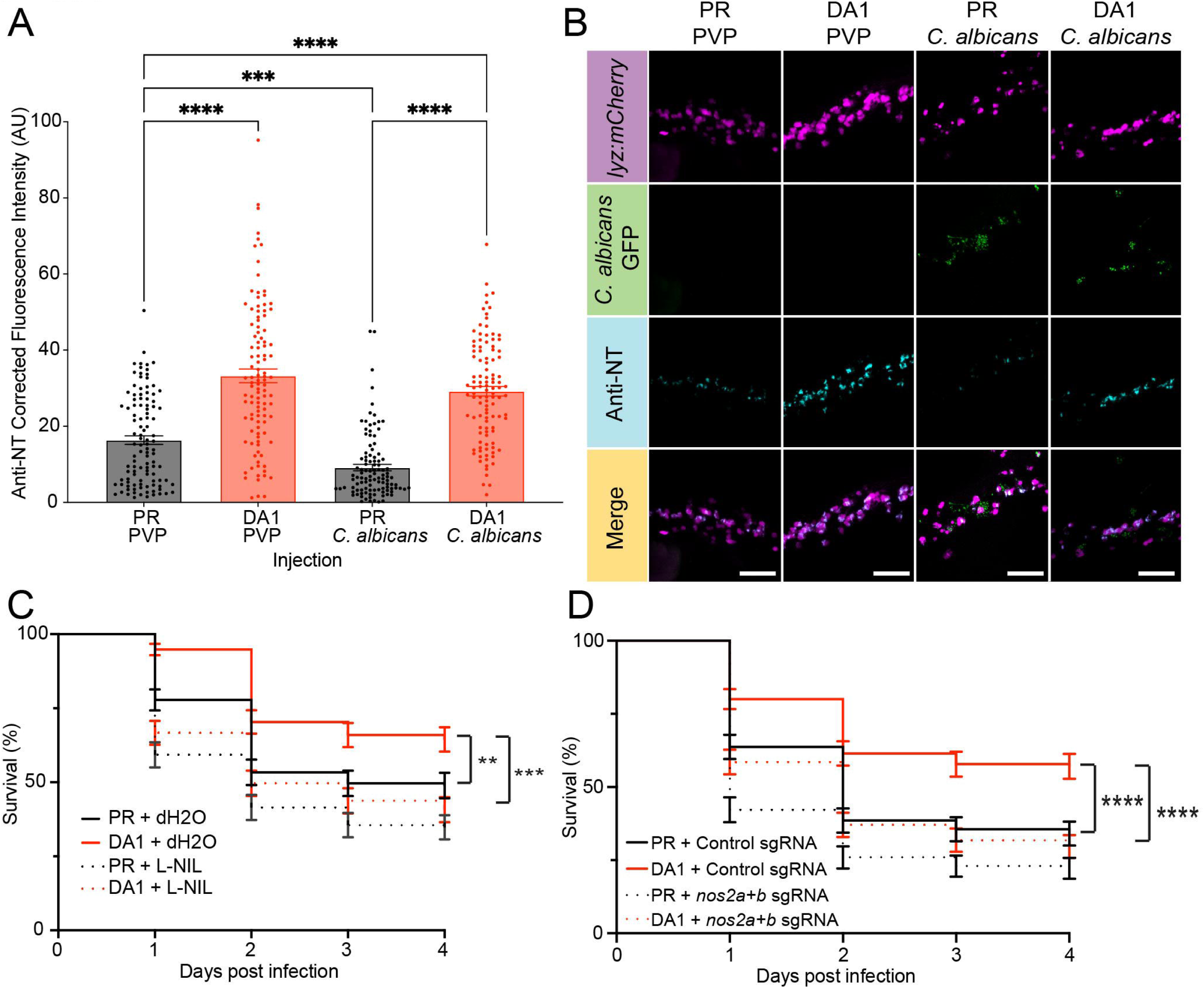
The host protective effect of Hif-1α stabilisation is RNS-dependent. (A) Anti-nitrotyrosine fluorescence at 1 dpi in PR and DA1 injected fish, following injection of PVP or 500 cfu *C. albicans* SN148 GFP. N=108 neutrophils from 18 fish, obtained from 3 independent experiments. Error bars show SEM. Statistical significance determined by Kruskal-Wallis test, with Dunn’s multiple comparisons test. ***p<0.001, ****p<0.0001 (B) Representative images of 1 dpi Tg(*lyz:NTRmCherry*) zebrafish larvae injected with PR or DA1 and infected with 500 cfu *C. albicans* SN148 GFP or PVP. Scale bars = 50 μm. (C) Survival curve of 200 cfu *C. albicans* TT21-dTomato infected zebrafish larvae following injection with PR or DA1, then treatment with dH2O or L-NIL. N=135 fish, obtained from 3 independent experiments. Error bars show SEM. Statistical significance determined by Gehan-Breslow-Wilcoxon test, with Bonferroni correction. **p<0.01, ***p<0.001. (D) Survival curve of 200 cfu *C. albicans* TT21-dTomato infected zebrafish larvae following injection with PR or DA1 and *tyr* or *nos2a+b* sgRNA. N=135 fish, obtained from 3 independent experiments. Error bars show SEM. Statistical significance determined by Gehan-Breslow-Wilcoxon test, with Bonferroni correction. **p<0.01, ***p<0.001.

To demonstrate that the increase in RNS after Hif-1α stabilisation is responsible for the host protective effect in *C. albicans* infection, we inhibited production of RNS both pharmacologically, using L-NIL, and genetically, using *nos2a+b* double CRISPR-Cas9 knockdown ("CRISPants"; Moore *et al*., 1994; Isles *et al*., 2019). L-NIL treatment did not impact the growth of *C. albicans in vitro* (Figure S8). Following infection with *C. albicans*, PR + L-NIL-treated larvae had reduced survival compared to PR + dH2O (drug solvent control)-treated larvae (Figure 7C), indicating a role of RNS in *C. albicans* control. DA *hif-1α* + L-NIL-treated larvae did not have a significant survival advantage compared to either PR + dH2O or PR + L-NIL groups (Figure 7C), demonstrating that iNOS inhibition is sufficient to remove the host protective effect of Hif-1α stabilisation. *Nos2a+b* double CRISPants were shown to inhibit nitrotyrosine production in mock infected zebrafish (Figure S9). DA *hif-1α* with *nos2a+b* CRISPants no longer had increased survival following *C. albicans* infection compared to PR + *tyr* sgRNA (*tyrosinase* CRISPant control; Isles *et al*., 2019) larvae, supporting that Nos2 is required for Hif-1α stabilisation dependent host protection (Figure 7D). Together these data demonstrate that the Hif-1α stabilisation is host-protective in *C. albicans* infection via a neutrophil- (Figure 6) and RNS-dependent (Figure 7) mechanism.

### Increased neutrophil RNS has an additive host protective effect when used alongside antifungals

We next investigated the potential of Hif-1α stabilisation to act as an adjunctive therapy alongside established antifungals. At 1 dpf, PR/DA *hif-1α* larvae were injected into the caudal vein with 500 cfu *C. albicans* TT21-dTomato then treated by immersion with 5.0 μg/ml fluconazole or 1.0 µg/ml caspofungin. Both fluconazole (Figure S10) and caspofungin (Figure S11) treatment had dose dependent effects on zebrafish larval survival post *C. albicans* infection. Fluconazole significantly increased host survival in both PR and DA *hif-1α* larvae (Figure 8A). DA *hif-1α* + fluconazole had increased survival compared to PR + fluconazole and DA *hif-1α* + DMSO. Combination of DA *hif-1α* + caspofungin similarly increased survival compared to PR + caspofungin (Figure 8B) and DA *hif-1α* + dH2O (Figure 8B), demonstrating that combination of Hif-1α stabilisation with fluconazole or caspofungin has an additive effect on survival.

**Figure 8:**
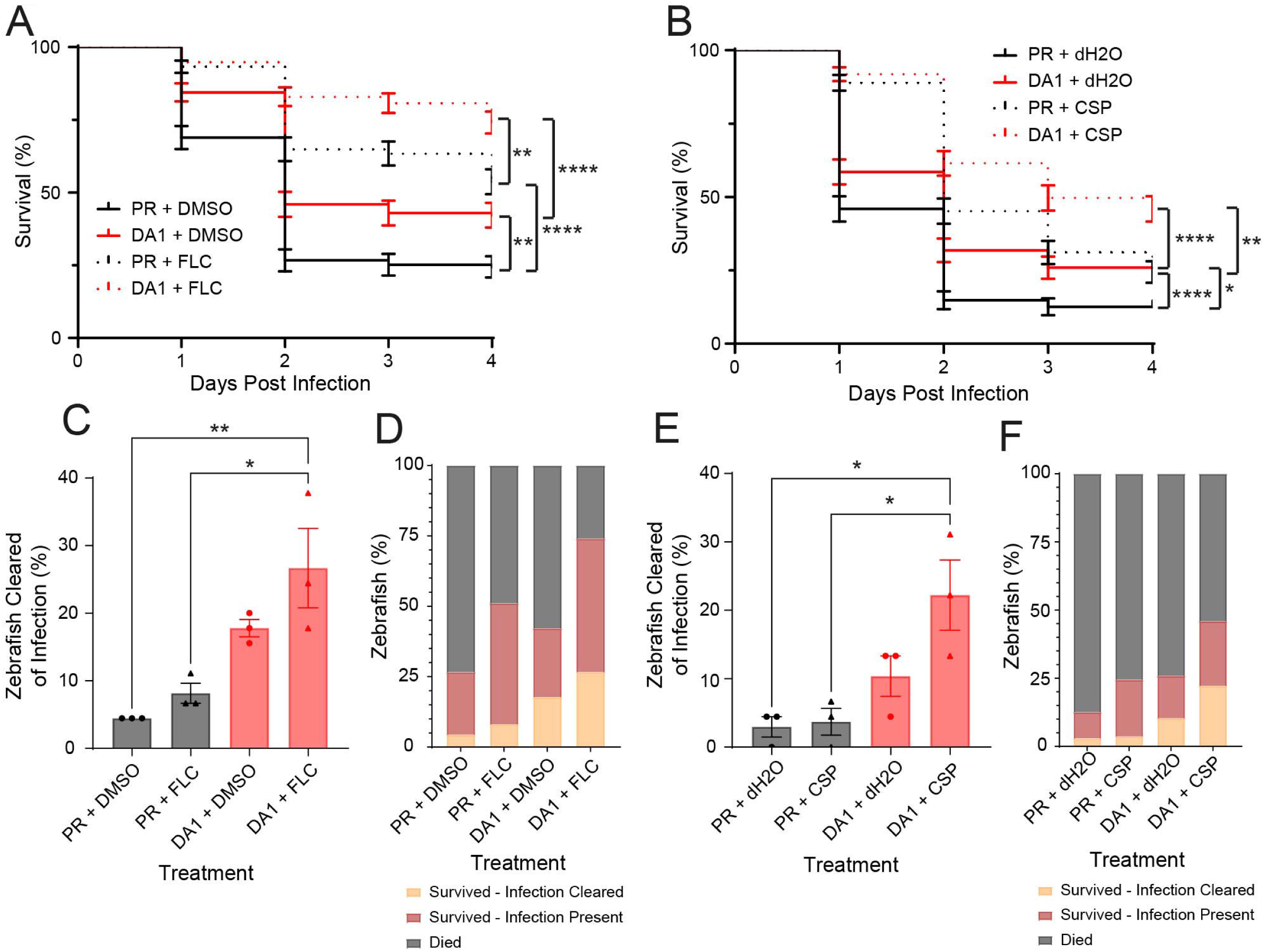
Hif-1α stabilisation has an additive effect with antifungals. (A) Survival of *C. albicans* TT21-dTomato infected zebrafish larvae following injection with dominant active *hif-1α* (DA1) or phenol red (PR) control. Larvae were treated with 5.0 µg/ml fluconazole or DMSO solvent control. Mortality was measured daily. N=135 fish, obtained from 3 independent experiments. Error bars show SEM. Statistical significance determined by Gehan-Breslow-Wilcoxon test with Bonferroni correction. **p<0.01, ****p<0.0001 (B) Survival of *C. albicans* TT21-dTomato infected zebrafish larvae following injection with DA1or PR control. Larvae were treated with 1.0 µg/ml caspofungin or dH2O solvent control. Mortality was measured daily. N=135 fish, obtained from 3 independent experiments. Error bars show SEM. Statistical significance determined by Gehan-Breslow-Wilcoxon test with Bonferroni correction. **p<0.01, ****p<0.0001 (C) Clearance of *C. albicans* infection at 4 dpi by zebrafish larvae following injection with DA1 or PR control and treatment with 5.0 µg/ml fluconazole or DMSO solvent control. N=3 independent experiments, 135 fish per group. Error bars show Standard Deviation. Statistical significance determined by one-way ANOVA, with Tukey’s multiple comparisons test. *p<0.05, **p<0.01 (D) Infection outcomes of *C. albicans* TT21-dTomato infected zebrafish larvae at 4 dpi, following injection with DA1 or PR and treatment with 5.0 µg/ml fluconazole or DMSO solvent control. N=3 independent experiments, 135 fish per group. (E) Clearance of *C. albicans* infection at 4 dpi by zebrafish larvae following injection with DA1 or PR control and treatment with 1.0 µg/ml caspofungin or dH2O solvent control. N=3 independent experiments, 135 fish per group. Error bars show Standard Deviation. Statistical significance determined by one-way ANOVA, with Tukey’s multiple comparisons test. (F) Infection outcomes of *C. albicans* TT21-dTomato infected zebrafish larvae at 4 dpi, following injection with DA1 or PR and treatment with 1.0 µg/ml caspofungin or dH2O solvent control. N=3 independent experiments, 135 fish per group.

Surviving larvae were imaged at 4 dpi to quantify clearance of *C. albicans* infection (Figure 8C-F). 27% of DA *hif-1α* + fluconazole larvae had completely cleared infection at 4 dpi (Figure 8C-D), greater than both PR alone (4%) and PR + fluconazole (8%). 22% of larvae treated with DA *hif-1α* + caspofungin cleared infection (Figure 8E-F) compared to 3% of PR-treated larvae and 4% of larvae treated with PR and caspofungin. Hence, Hif-1α stabilisation also has an additive effect on clearance when combined with fluconazole or caspofungin.

## Discussion

The increasing prevalence of antifungal resistance worldwide highlights the importance of discovering new antifungal targets or host-directed therapies capable of improving disease outcomes. Here, we have shown that *Candida* spp. suppression of neutrophil RNS is an important factor in their virulence *in vivo* and that restoration of neutrophil RNS represents a potential host directed therapy for difficult-to-treat *C. albicans* infection.

We found that *C. albicans* can suppress neutrophil RNS production in zebrafish *in vivo* and in human primary neutrophils *in vitro*. *C. albicans* has been shown to suppress RNS production in murine bone marrow derived macrophages *in vitro* (Collette, Zhou and Lorenz, 2014), which is supported by a variety of sources demonstrating suppression of ROS by *C. albicans in vitro* (Frohner *et al*., 2009; Wellington, Dolan and Krysan, 2009). While previous publications have demonstrated RNS suppression in macrophages, our data demonstrates an effect in neutrophils in a live tissue-setting. Human alveolar macrophages are poor RNS producers (Muijsers *et al*., 2001) and *C. albicans* is primarily controlled by neutrophils in human disease (Gazendam *et al*., 2016), therefore we chose to dissect the mechanisms behind RNS suppression in neutrophils in the tractable *in vivo* zebrafish model.

Here, we identify that RNS suppression is partially dependent on live, hyphae-forming *C. albicans*. The hyphal morphotype is highly associated with a shift in transcription and increased virulence (Sudbery, 2011; Desai, 2018), which may trigger expression of protein(s) necessary for RNS suppression by *C. albicans*. Proteomic analysis of supernatant from hyphal *C. albicans in vitro* identified 301 secreted proteins specific to hyphae (Vaz *et al*., 2021).

Further examination and characterisation of these proteins could point towards candidates involved in active RNS suppression. However, RNS suppression was still observed in non-hyphae-forming, non-albicans *Candida* spp. isolates (*C. glabrata, C. parapsilosis, C. guilliermondii*), suggesting that the presence of hyphae themselves is not essential for RNS suppression.

Host and *C. albicans* derived arginase also contribute to the suppression of neutrophil RNS production. CAR1 is the only known arginase gene in the *C. albicans* genome, encoding a cytosolic arginase (Schaefer *et al*., 2020). Fungal arginase may be interfering with the host arginase-iNOS balance, resulting in reduced iNOS activity and RNS suppression. The modest decrease in neutrophil RNS production after infection with the CAR1 mutant suggests that *C. albicans* arginase production is not the sole RNS reducing mechanism. Our findings suggest that multiple, additive mechanisms of neutrophil RNS suppression by *C. albicans* may underlie their success at the large overall suppression of neutrophil RNS observed. *C. albicans* SOD5 has been associated with degradation of antimicrobial ROS *in vitro* (Frohner *et al*., 2009), *C. albicans* chitin induces host arginase-1 activity, decreasing RNS production (Wagener *et al*., 2017), *YHB1* is able to detoxify RNS (Hromatka, Noble and Johnson, 2005) and Collette *et al*. identified an unknown, small secreted molecule capable of suppressing RNS (Collette, Zhou and Lorenz, 2014). The presence of chitin may account for the partial suppression of RNS by heat-killed *C. albicans* that we observed. Further investigation could aim to establish which are the most important mechanisms and effector proteins involved in RNS suppression in neutrophils.

Neutrophil RNS suppression was observed in infection with various *C. albicans* laboratory strains and clinical isolates alongside non-albicans *Candida* spp. clinical isolates. This suggests that RNS suppression could be conserved widely across clinically relevant *Candida* spp. Importantly, *Candida* spp. virulence correlated with neutrophil RNS suppression *in vivo* indicating that neutrophil RNS suppression may be a virulence factor. Virulence is determined by a wide range of factors, not solely neutrophil RNS suppression (Ghannoum and Abu-Elteen, 1990), any of which could be differentially regulated in these *Candida* spp. isolates. It was interesting to note that the emerging human pathogen *C. auris* was able to efficiently suppress neutrophil RNS in zebrafish, despite only been identified as a human pathogen since 2009 (Lockhart *et al*., 2017), suggesting that this immune evasion mechanism may have evolved in the environment (Casadevall, Kontoyiannis and Robert, 2019). Suppression of neutrophil RNS by environmental strains may act as a useful proxy for determining whether *Candida* spp. or strains have the potential to be virulent in humans.

We found that upregulation of RNS, via Hif-1α stabilisation, has a protective effect in *C. albicans* infection in zebrafish larvae, via a neutrophil-mediated, RNS-dependent mechanism. Neutrophils are the primary innate immune cell responsible for killing *Candida spp.* (Brown, 2011). Neutrophil suppression is therefore an important mechanism of *C. albicans* pathogenesis that, if restored, could represent an exciting therapeutic opportunity. RNS is well characterised as being candidacidal (Rementería, García-Tobalina and Sevilla, 1995; Vazquez-Torres *et al*., 1995; Tillmann, Gow and Brown, 2011; Navarathna, Lionakis and Roberts, 2019) and NO-releasing nanoparticles inhibited the growth of *C. albicans in vitro* and *in vivo* (Macherla *et al*., 2012). Hence, upregulation of neutrophil RNS production is a potential mechanism that could be targeted therapeutically to improve immune candidacidal activity. We have previously demonstrated Hif-1α stabilisation protects against *M. marinum* infection in a similar manner (Elks *et al*., 2013), suggesting Hif-1α stabilisation could offer protective effect against multiple and different groups of pathogens (bacteria and fungi) where neutrophil-mediated immune responses are dampened during pathogenesis. Further to this, we showed restoration of neutrophil RNS had an additive effect on survival and clearance when combined with traditional antifungals. Early fungicidal activity by neutrophils prior to hyphal growth is vital for subsequent clearance and resolution of *C. albicans* infection, with early phagocytosis a strong prognostic indicator of survival (Bergeron *et al*., 2017; Zhu *et al*., 2023). As a primarily fungistatic agent (Vasicek *et al*., 2014), fluconazole may help to restrict initial growth of *C. albicans*, allowing neutrophils with increased RNS production to kill *C. albicans,* resulting in greater survival and clearance. Conversely, caspofungin predominantly has a fungicidal effect (Letscher-Bru and Herbrecht, 2003; Szymański *et al*., 2022), though increased neutrophil RNS production may also facilitate earlier killing of *C. albicans* in combination and allow lower doses of these poorly tolerated drugs for shorter treatment periods.

Our data shows that *Candida* spp. can efficiently suppress neutrophil RNS as a virulence mechanism and that Hif-1α stabilisation is able to overcome this, conferring a neutrophil-mediated, RNS-dependent protective effect *in vivo*. Neutrophil RNS restoration has the potential as an adjunctive host-directed therapeutic strategy to aid in the treatment of *Candida* spp. infections.

## Acknowledgements

The authors would like to thank The BSU Aquarium Team for fish care and the SMPH Technical Team for practical assistance (University of Sheffield). We would like to thank Professor Neil Gow (University of Exeter), Dr Robert Wheeler (University of Maine) and Dr Ben Caswall (St George’s, University of London) for kindly providing *Candida* spp. strains. For the purpose of open access, the author has applied a Creative Commons Attribution (CC BY) licence to any Author Accepted Manuscript version arising

## Author Contributions

Conceived and designed the experiments: TBB, FRH, PTS, AL, SC, SAJ, AMC, PME. Performed the experiments: TBB, FRH, PTS, AL. Generation/provision of *Candida spp*. strains: SC, TB KRA, DGP. Analysed the data: TBB, FRH, PTS, AL, AMC, PME. Wrote and drafted the manuscript: TBB, FRH, PTS, AL, SAJ, AMC, PME.

