## Supplemental Figures for "*Candida spp.* suppress neutrophil reactive nitrogen species to evade killing"

### Supplementary Figures and Legends

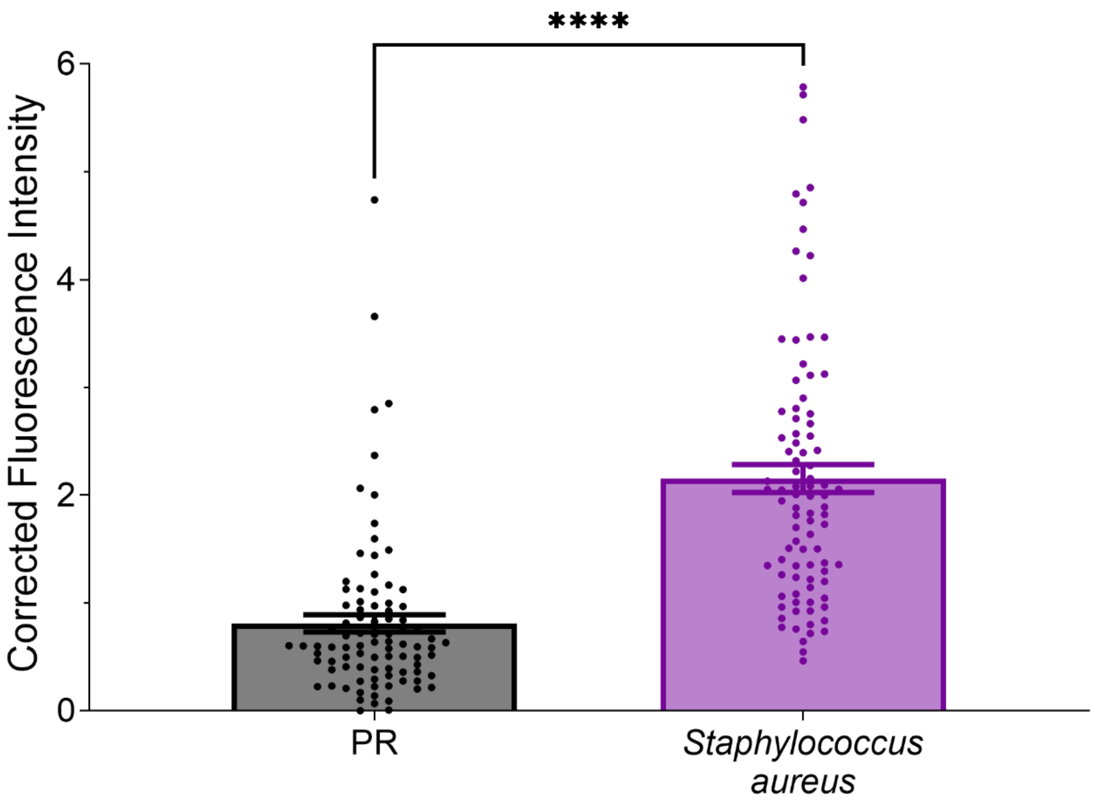


**Figure S1: *S. aureus* increases neutrophil RNS *in vivo*.**

Anti-nitrotyrosine fluorescence at 1 dpi following injection of PR or *S. aureus* (SH1000) into the caudal vein. N=90 neutrophils from 15 fish, obtained from 3 independent experiments. Error bars show SEM. Statistical significance determined by two-tailed Mann-Whitney test. P values shown: ****$p<0.0001$.

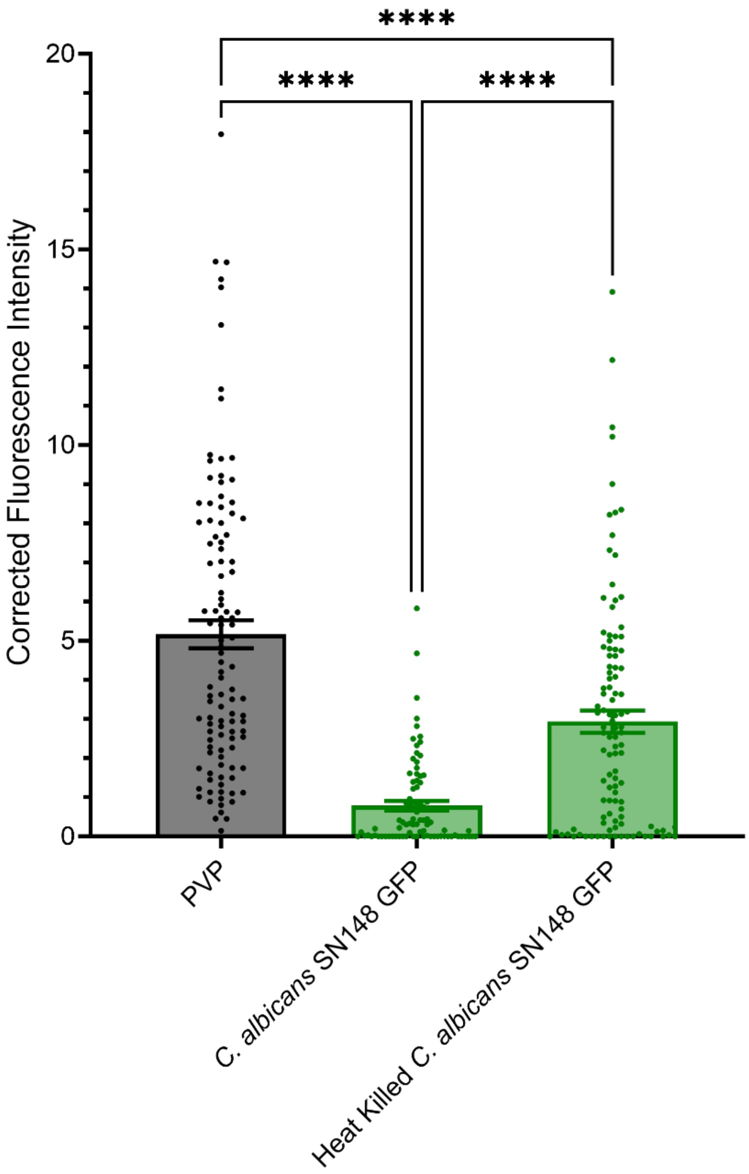


**Figure S2: *C. albicans* SN148 GFP suppresses neutrophil RNS.**

Anti-nitrotyrosine fluorescence at 24 hpi following injection of PVP, *C. albicans* TT21 dTomato or heat killed *C. albicans* TT21 dTomato into the caudal vein. N=84-108 neutrophils from 14-18 fish, obtained from 3 independent experiments. Error bars show SEM. Statistical significance determined by Kruskal-Wallis test, then Dunn's multiple comparisons test. ****p<0.0001.

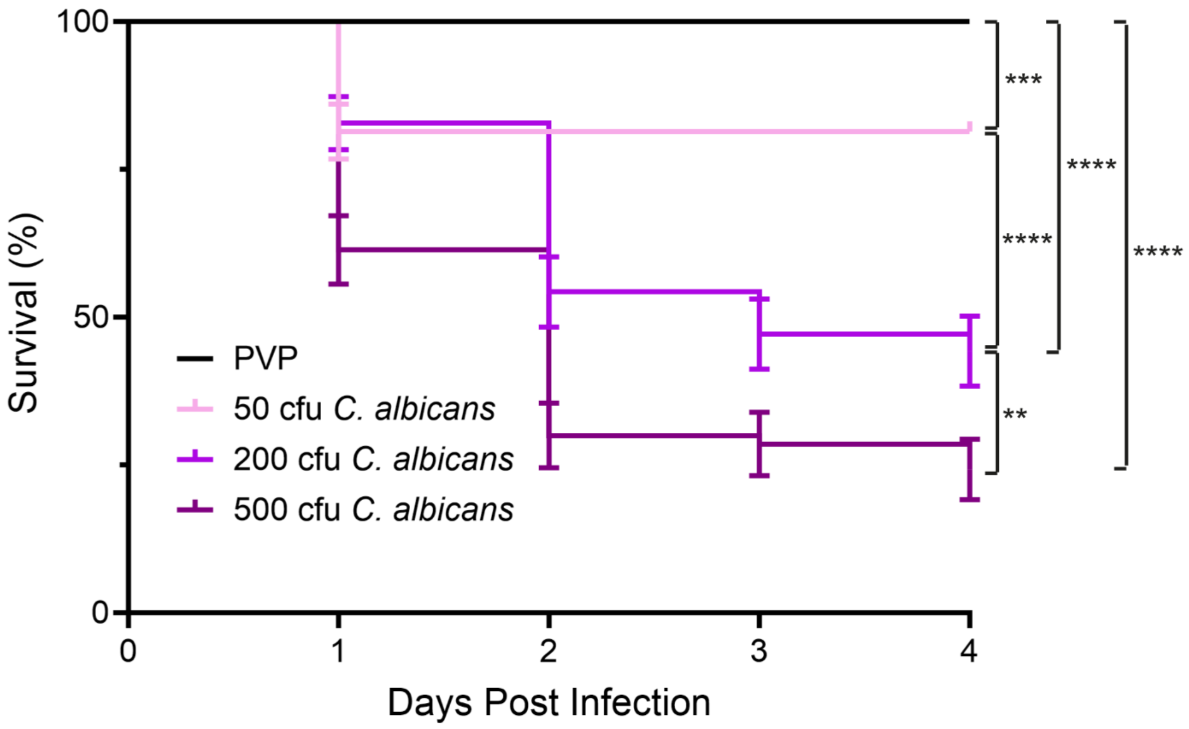


**Figure S3: *C. albicans* has a dose-dependent effect on survival.**

Survival curve of PVP, 50 cfu *C. albicans*, 200 cfu *C. albicans* and 500 cfu *C. albicans* infected zebrafish larvae. Mortality was measured daily. N=70 fish, obtained from 2 independent experiments. Error bars show SEM. Statistical significance determined by Gehan-Breslow-Wilcoxon test, with Bonferroni correction. **$p<0.01$, ***$p<0.001$, ****$p<0.0001$.

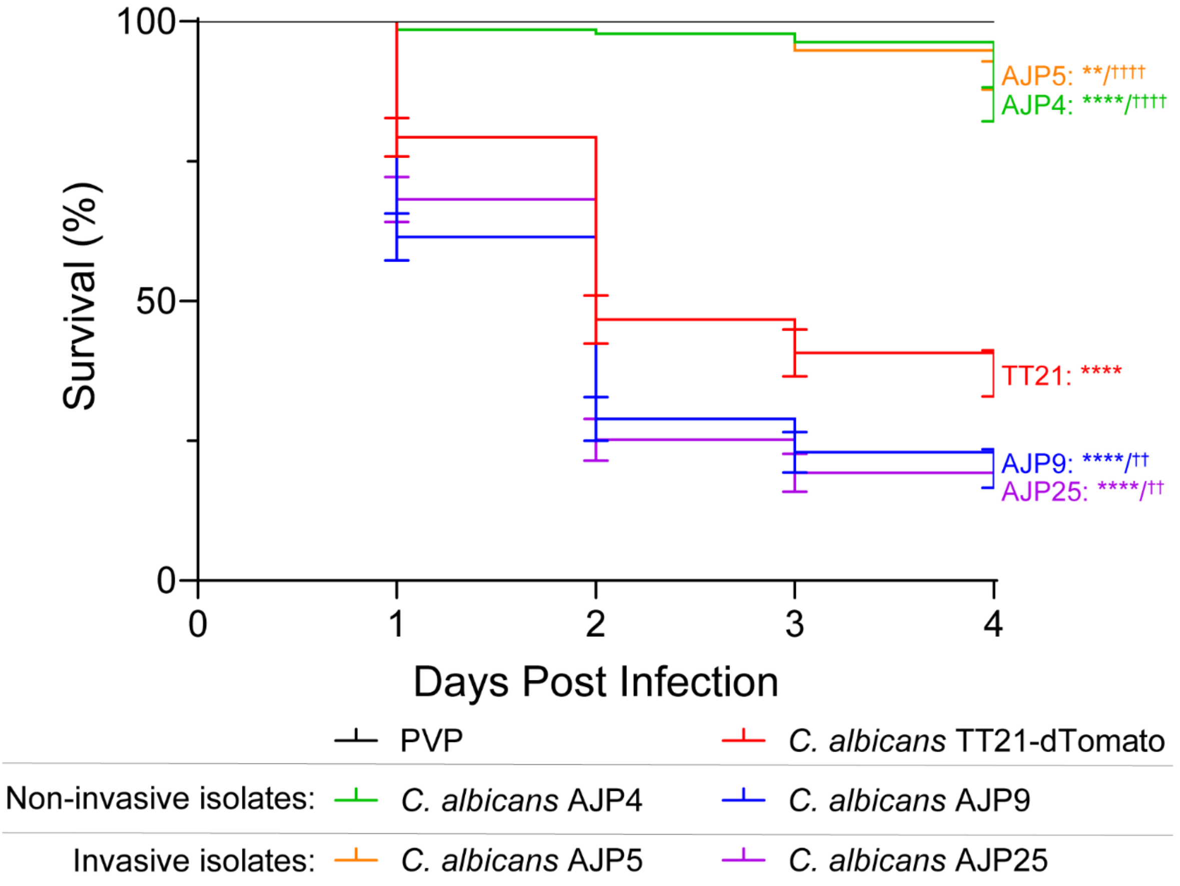


**Figure S4: *C. albicans* clinical isolates have varying levels of virulence in zebrafish.** 1 dpf nacre embryos were injected into the caudal vein with PVP, 200 cfu *C. albicans* TT21-dTomato, AJP4, AJP5, AJP9, AJP25 or PVP. Mortality was measured daily up to 4 dpi. n=135 fish, obtained from 3 independent experiments. Statistical significance determined by Gehan-Breslow-Wilcoxon test, with Bonferroni correction. * indicates difference compared to PVP. † indicates difference compared to *C. albicans* TT21-dTomato. P values shown: **/††p<0.01, ****/††††p<0.0001.

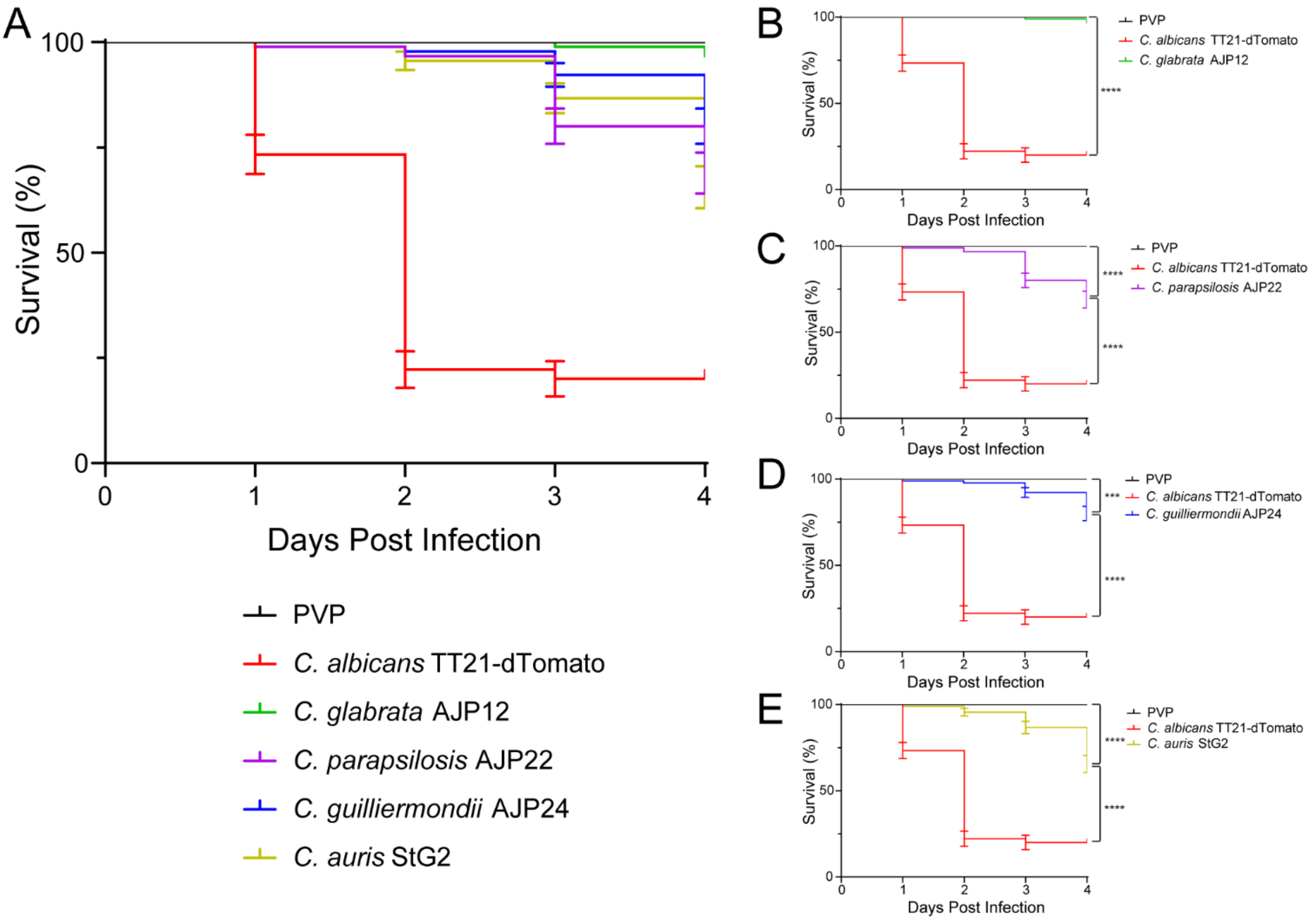


**Figure S5: Non-albicans *Candida* spp. clinical isolates have varying levels of virulence in zebrafish.**

(A) 1 dpf nacre embryos were injected into the caudal vein with 200 cfu *C. albicans* TT21-dTomato, *C. glabrata* AJP12, *C. parapsilosis* AJP22, *C. guilliermondii* AJP24, *C. auris* StG2 or PVP. Mortality was measured daily up to 4 dpi. n=90 fish, obtained from 2 independent experiments. Statistical significance determined by Gehan-Breslow-Wilcoxon test, with Bonferroni correction. Statistical differences only shown on (B-E) for simplicity.

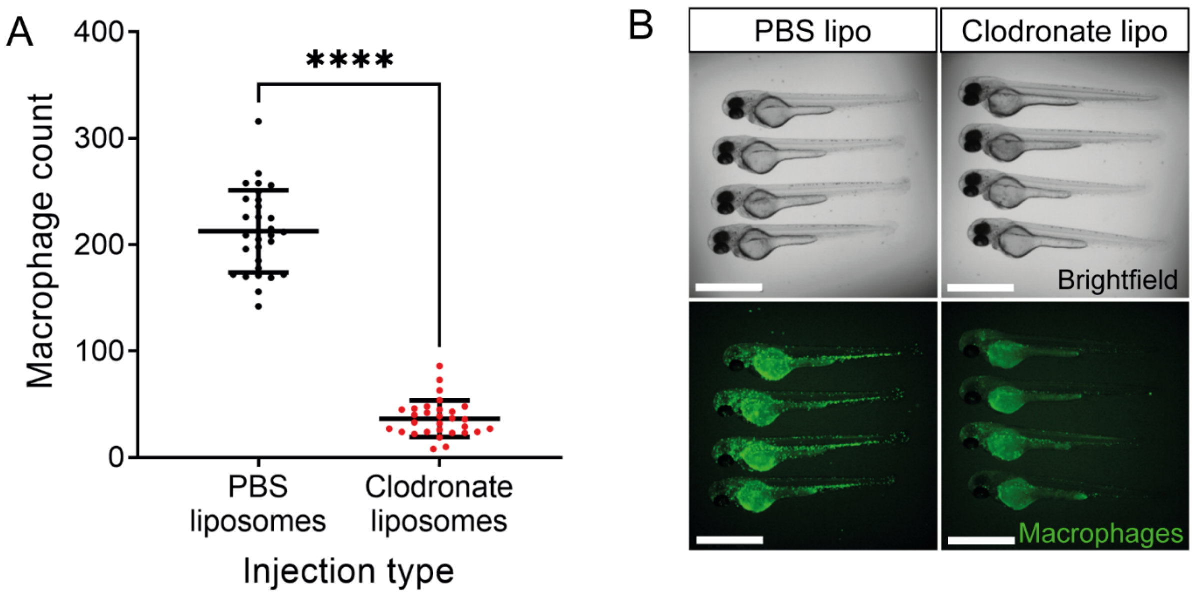


**Figure S6: Clodronate liposomes deplete the macrophage population.**

(A) Total body macrophage count at 2 dpf (1 dpi). N=30 fish, obtained from 3 independent experiments. Error bars show SEM. Statistical significance determined by unpaired T test. ****p<0.0001.

(B) Representative images of 2 dpf (1 dpi) Tg(*mpeg:nlsClover*) zebrafish embryos following injection with PBS or clodronate liposomes.

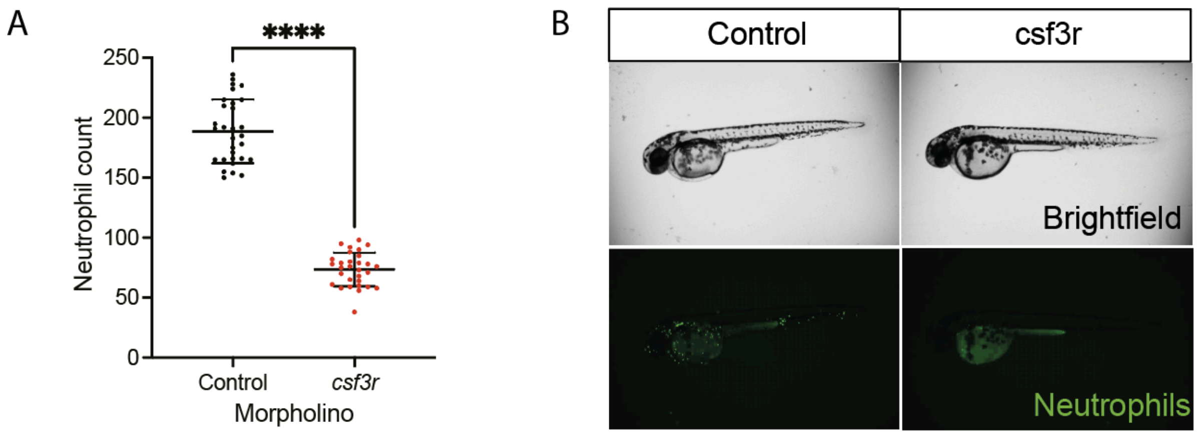


**Figure S7: *csf3r* morpholino depletes the neutrophil population.**

(A) Total body neutrophil count at 2 dpf. N=30 fish, obtained from 3 independent experiments. Error bars show SEM. Statistical significance determined by unpaired T test. ****p<0.0001.

(B) Representative images of 2 dpf Tg(*mpx:GFP*) i114 zebrafish embryos following injection with Control or *csf3r* morpholino.

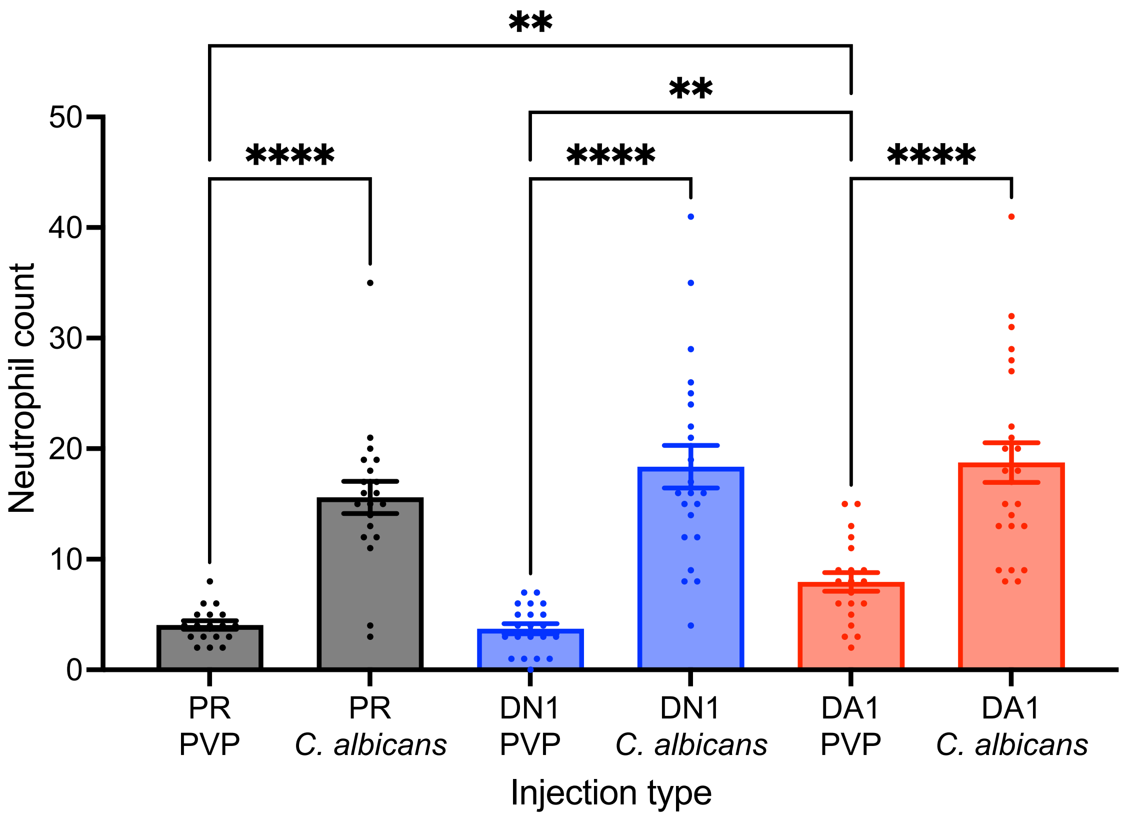


**Figure S8: DA1 does not increase recruitment of neutrophils to *C. albicans* infection.** Number of neutrophils recruited to site of injection at 24 hpi, following injection with PVP or 20 cfu *C. albicans* TT21 dTomato into the hindbrain ventricle. Embryos were previously injected with PR, DN1 or DA1. N=18-24 fish, obtained from 3 independent experiments. Statistical significance determined by Brown-Forsythe and Welch ANOVA with Dunnett's T3 multiple comparisons test. **p<0.01, ****p<0.0001.

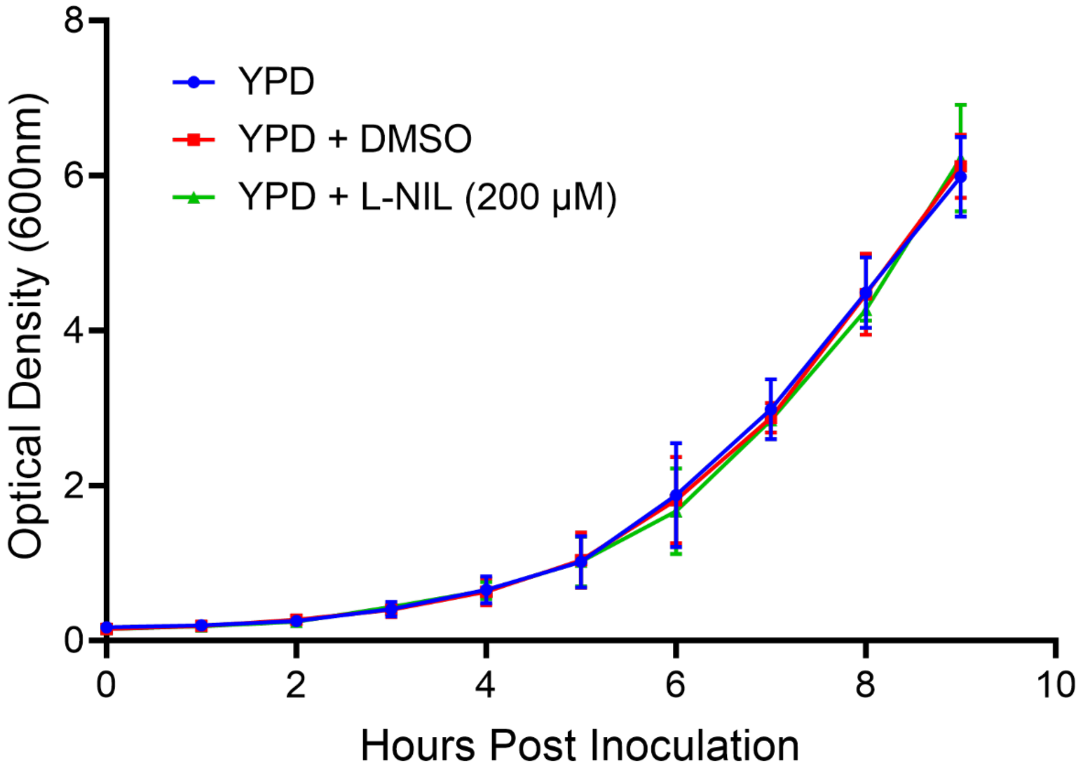


**Figure S9: L-NIL does not affect growth of *C. albicans in vitro*.**

Growth curve of *C. albicans* TT21 dTomato grown in YPD, YPD + DMSO or YPD + L-NIL over 10 hours. Optical Density (600nm) was measured every hour. Error bars show SD. Comparison of Fits finds no statistically significant difference between groups ($p=0.9929$), based on exponential growth model.

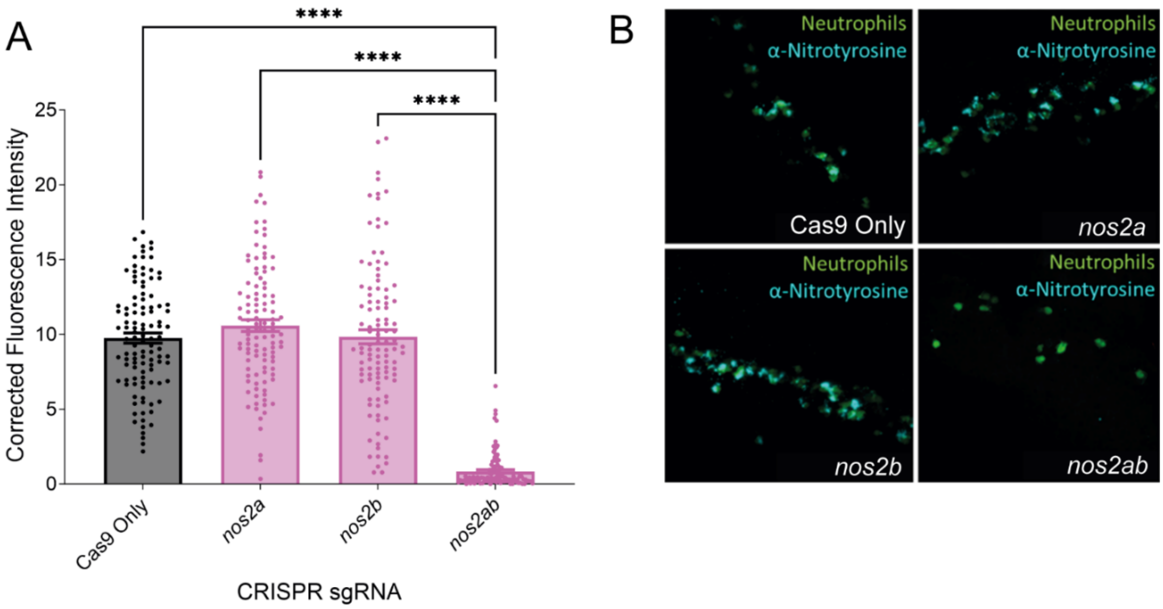


**Figure S10: *nos2ab* double CRISPR prevents production of RNS.**

(A) Anti-nitrotyrosine fluorescence in 5 dpf zebrafish embryos following injection of Cas9 only, *nos2a* sgRNA, *nos2b* sgRNA or *nos2ab* sgRNA. Error bars show SEM. N=108 neutrophils from 18 fish, obtained from 3 independent experiments. Statistical significance determined by Kruskal-Wallis test, with Dunn's multiple comparisons test. ****$p<0.0001$ (B) Representative images of nitrotyrosine stained zebrafish embryos.

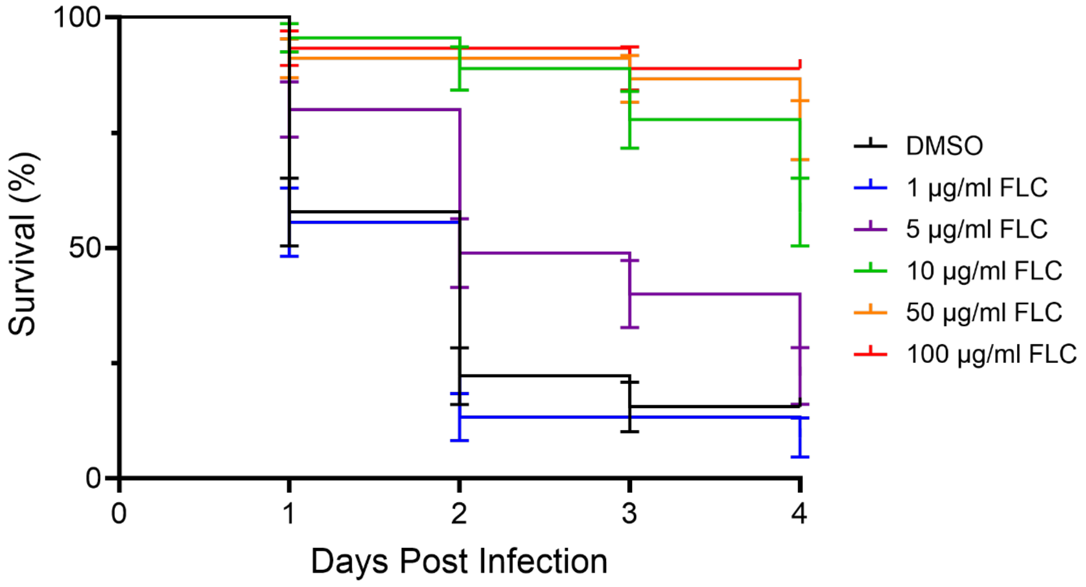


**Fig S11: Fluconazole has a dose dependent effect on survival in *C. albicans* infection.** Survival of 500 cfu *C. albicans* TT21-dTomato infected zebrafish embryos treated with a range of fluconazole concentrations or DMSO vehicle control. N=45 fish, obtained from 1 independent experiment. Error bars show SEM. No statistical tests were performed.

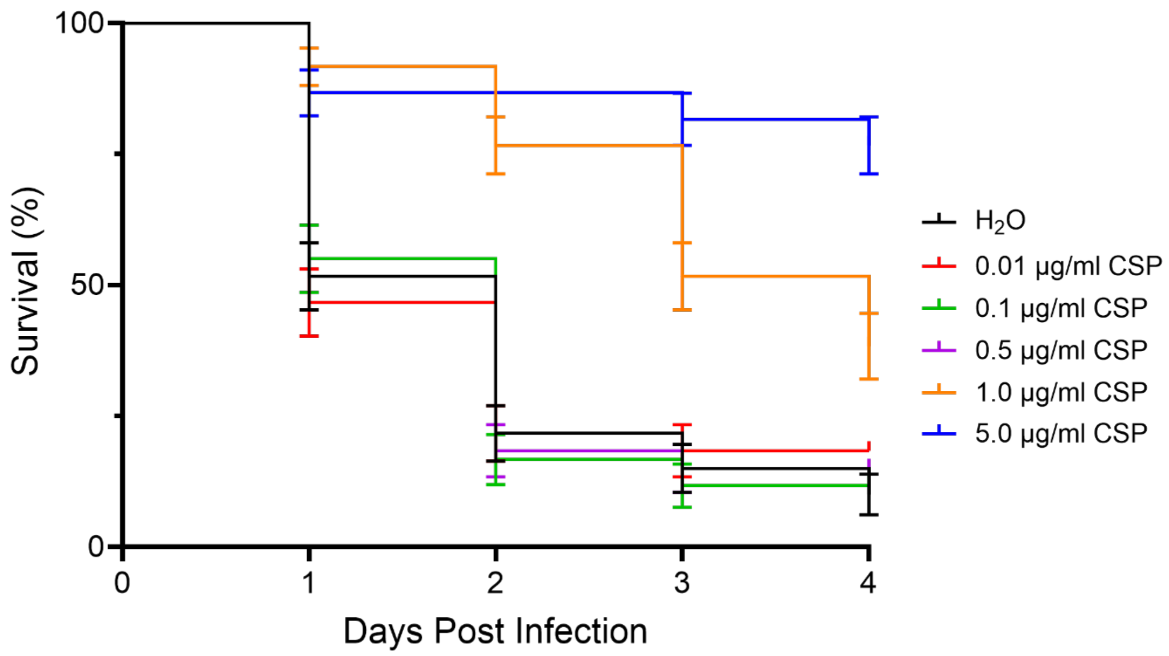


**Figure S12: Caspofungin has a dose dependent effect on survival in *C. albicans* infection.**

Survival of 500 cfu *C. albicans* TT21-dTomato infected zebrafish embryos treated with a range of caspofungin concentrations or d$H_2O$ solvent control. N=60 fish, obtained from 1 independent experiment. Error bars show SEM. No statistical tests were performed.
